# Bacterial glycosphingolipids orchestrate colonization and immune modulation in neonatal host

**DOI:** 10.1101/2025.05.02.651985

**Authors:** Kyoo Heo, Da-Jung Jung, Ji-Sun Yoo, Byoungsook Goh, Dennis L. Kasper, Sungwhan F. Oh

**Affiliations:** Center for Experimental Therapeutics and Reperfusion Injury, Department of Anesthesiology, Perioperative and Pain Medicine, Brigham and Women s Hospital, Boston, MA, USA; Department of Immunology, Blavatnik Institute of Harvard Medical School, Boston, USA; Graduate Program in Immunology, Division of Medical Science, Harvard Medical School, Boston, USA

## Abstract

Symbiotic microbiota has co-existed with the mammalian host over millennia, conserving a stable community structure generation after generation. During the vertical transmission, gut symbionts rapidly colonize the unoccupied host lumen, nonetheless, how symbionts adapt to the dynamic changes of host environment, and contribute to the structural and immunological maturation remains elusive. Here, we show that the early gut symbiont *Bacteroides fragilis* produces a species- and stage-specific sphingolipid, alpha-galactosylceramide (BfaGC), that orchestrates neonatal colonization and immune modulation. BfaGC stabilizes membrane integrity and facilitates aerobic respiration, providing a critical advantage under early-life oxygen exposure. Temporally induced in the neonatal gut, BfaGC is necessary to regulate colonic type I natural killer T (NKT) cells, highlighting metabolic adaptation of the symbiont is synchronized with the time-sensitive host development. These findings reveal a mutualistic benefit exerted by endobiotic lipid metabolites in host-microbe interactions and provide new insights into species-specific mechanisms in early microbiota establishment and host immune education.

## Main

The human gut microbiota exerts profound influence on host physiology, shaping nutrient metabolism, immune system development, and overall intestinal homeostasis^1-4^. The neonatal stage is a time of dynamic change for the initially introduced microbial community, including transition in nutrient availability and immune maturation^5-7^. Adapting to both host and environmental factors, gut microbiota undergo rapid assembly, establishing a foundation that can influence lifelong health outcomes^8,9^. Deviations in such process (e.g. non-vaginal delivery, antibiotics use, and unconventional feeding practices) are associated with elevated risks of allergies, autoimmune disorders, and metabolic diseases^10^, underscoring the far-reaching consequence of neonatal microbial colonization.

Despite the recognized importance of early microbial assembly, the mechanistic basis by which specific bacterial species thrive under neonatal conditions remains elusive. The neonatal gut differs markedly from that of adult’s: it is not yet strictly anaerobic, the immune system is still maturing, and the repertoire of available nutritional substrates can shift dramatically over the first few weeks of life in mouse, or first years in human^11,12^. Microbes must adapt to establish a foothold in this dynamic environment. Previous studies highlight certain taxa, such as *Bifidobacterium* and *Bacteroides*, as prominent early colonizers^13,14^, but why these organisms succeed where others fail has remained unclear.

Here, we address these gaps in understanding by systematically interrogating the genetic determinants of neonatal gut colonization by *Bacteroides fragilis* using genomewide screening of fitness in adult and neonatal guts. Our findings reveal that *B. fragilis* leverages specialized metabolic and stress-response pathways to thrive in the microaerobic neonatal gut^15^. Central to this adaptation is the production of alpha-galactosylceramide (aGC), which stabilizes membrane potential and enhances respiration under transient oxygen conditions, thereby conferring a pivotal advantage during early colonization and species-specific temporal niche acquisition. Additionally, these observations underscore a co-evolutionary dynamic wherein the metabolic function of BfaGC may have converged with host immune recognition within this critical neonatal time window, further solidifying *B. fragilis* as a favored early colonizer. By elucidating how glycolipid-mediated membrane adaptations enhance microbial fitness in the neonatal gut, our study reveals a fundamental mechanism by which metabolic flexibility shapes early colonization dynamics, providing a broader framework for understanding the ecological and evolutionary forces governing host-associated microbial communities.

## Results

### Genetic determinants of bacterial colonization in the neonatal gut

Longitudinal analysis of the human infant microbiome revealed that *B. fragilis* exhibits a significantly higher abundance in the neonatal gut compared to other *Bacteroidetes* species^14^. To recapitulate this phenomenon, we generated gnotobiotic mice colonized with a defined 10-member *Bacteroidetes* community (Fig. 1a). In the pregnant dam, *B. fragilis* abundance dropped to ∼2% and stabilized. In contrast, the gut microbiota of offspring showed a strong but transient domination of *B. fragilis* in early life (∼39.2% at postnatal day 14 (p14)). These findings suggest that *B. fragilis* employs unique colonization strategies in the neonatal gut, which remain effective in the murine model.

**Figure 1.**
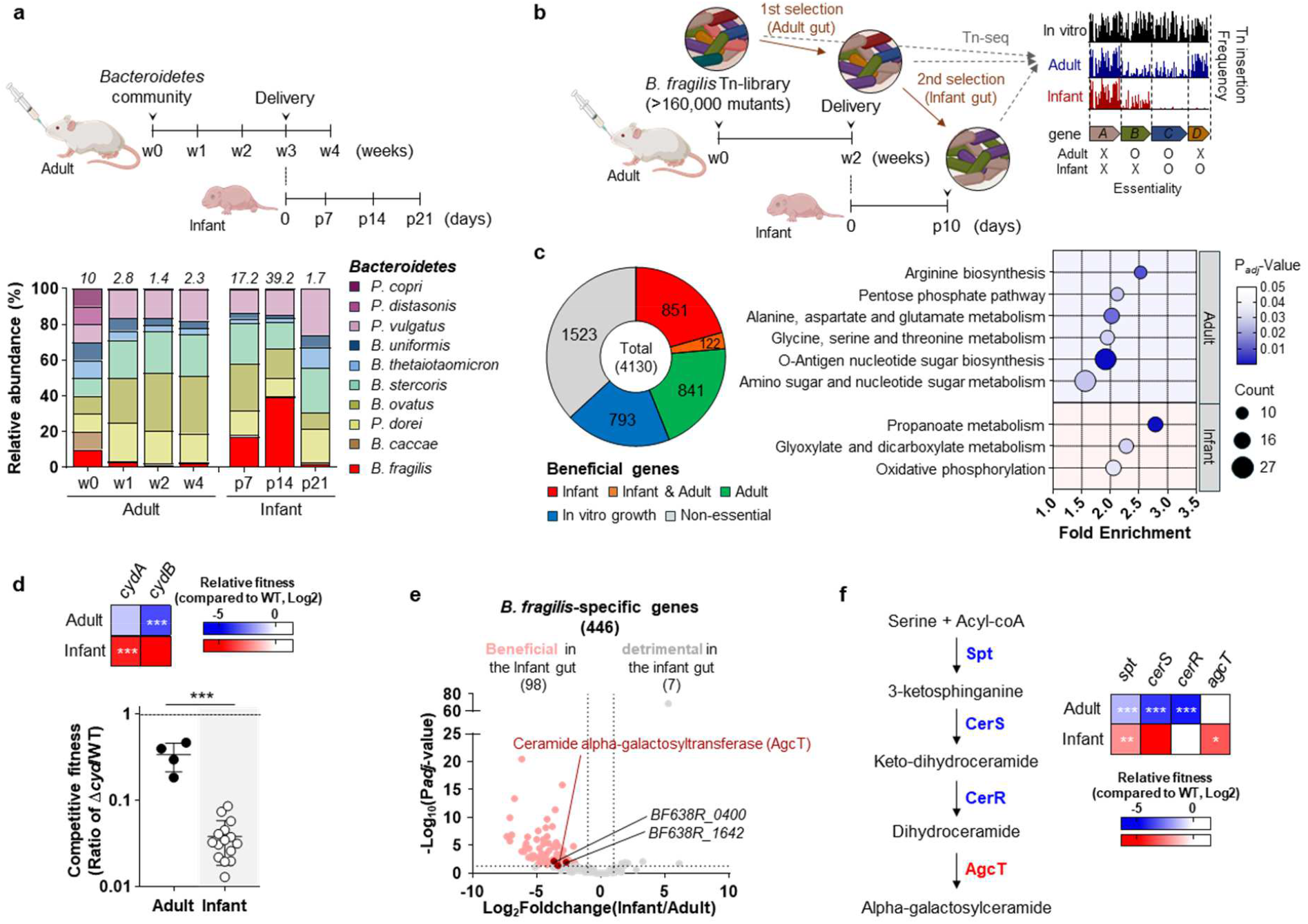
Genetic determinants of *Bacteroides fragilis* colonization in the neonatal gut. **a**, Longitudinal relative abundance of a defined consortium of 10 *Bacteroidetes* species (*B. fragilis, Bacteroides caccae, Bacteroides thetaiotaomicron, Bacteroides uniformis, Bacteroides stercoris, Phocaeicola vulgatus, Phocaeicola dorei, Bacteroides ovatus, Prevotella copri, Parabacteroides distasonis*), colonized in germ-free Swiss-webster mice. *Bacteroides fragilis* abundance is indicated in the graph. Data represent mean of n=6-7 mice per time point. **b**, Transposon sequencing (Tn-seq) workflow for identifying genes essential for colonization. A high-density Tn-mutant library of *B. fragilis* (∼160,000 mutants) was introduced into germ-free mice. The initial mutant pool and populations recovered from adult and infant mice were then analyzed via insertion sequencing. Genes with significantly decreased insertion frequency under a given condition were classified as beneficial for colonization in that environment. **c**, Classification of beneficial genes based on Tn-seq revealed condition-specific requirements for *B. fragilis* colonization in the adult and neonatal gut (left), with subsequent KEGG pathway enrichment analysis highlighting distinct metabolic and stress-response pathways engaged at each developmental stage (right). **d**, Identification of *cydAB*, encoding cytochrome *bd* oxidase, as a key determinant of *B. fragilis* survival in the neonatal gut, further validated by competitive colonization between wild-type and its isogenic Δ*cyd* mutant *B. fragilis* in neonatal and adult mice. **e**, Among *B. fragilis*-specific genes, those beneficial in the infant gut are indicated in pale red, and a subset previously shown to be oxygen-induced is marked in deep brown. *agcT*, encoding ceramide α-galactosyltransferase, is one of only three genes belonging to both categories, uniquely positioning it as a key oxygen-responsive determinant of neonatal colonization. **f**, Relative fitness of transposon-insertion mutant of core ceramide synthesis genes (*spt, cerR, cerS*) and *agcT* was determined compared to WT in both adult and infant gut. Shown are the means and SD.

Previous studies have implicated several genetic traits in facilitating *Bacteroidetes* colonization of the neonatal gut^16^. To systematically define the genetic basis of early-life colonization, we generated a high-density *B. fragilis* mutant library (>160,000 isogenic mutants) and performed transposon sequencing (Tn-seq) to identify beneficial genetic factors **(Fig. 1b and Fig. S1a)**^17,18^. The library was introduced into pregnant germ-free mice, allowing natural vertical transmission to neonates and stage-specific *in vivo* selection. Prior to *in vivo* selection, 80.8% of open reading frames (ORFs) harbored sufficient transposon insertions, indicating that most genes are non-essential under nutrient-rich conditions **(Fig. S1b)**. In contrast, mutant diversity decreased markedly *in vivo*—from 160,965 unique insertion sites *in vitro* to 43,816 in adults and 12,110 in neonates **(Fig. S1c)**. To distinguish genuine selection from stochastic loss due to in vivo bottlenecking, we applied a zero-inflated negative binomial (ZINB) model that accounts for sparse and zero-heavy data, enabling robust identification of genes significantly depleted in neonates. Using this framework, we identified 963 and 973 genes contributing to *B. fragilis* survival in the adult and infant gut, respectively **(Fig. 1c, Fig. S1d, and Supplementary Table 1)**. In adults, enriched pathways included carbohydrate and amino acid metabolism, suggesting nutrient independence as a key fitness trait, while O-antigen biosynthesis, especially the synthesis of polysaccharide A (PSA), indicated dual roles for extracellular polysaccharides in both microbial survival and host interaction **(Fig. 1c, Supplementary Table 2, and Fig. S2)**^19^.

In neonates, colonization relied on distinct metabolic pathways, with propanoate metabolism, glyoxylate/dicarboxylate metabolism, and oxidative phosphorylation **(Fig. 1c)**. Among the enriched pathways, oxidative phosphorylation was of particular interest given its role in aerobic respiration, reflected in the essentiality of *cydAB* (cytochrome *bd* oxidase) in the neonatal gut **(Fig. 1d)**^20,21^. Its deletion led to a marked colonization defect (Competitive index = 0.038 in the neonates, 0.337 in the adult), revealing that *B. fragilis*, although a strict anaerobe, engages in oxygen-dependent metabolism during early colonization. While cytochrome *bd*-mediated respiration facilitates bacterial adaptation near the gut epithelium in the adult gut^22,23^, its essentiality in neonates underscores a heightened reliance on this pathway before strict anaerobiosis is fully established^15^. The conservation of *cydAB* across *Bacteroides* species supports its importance as a core adaptation to fluctuating oxygen environments^21^.

These respiratory adaptations were accompanied by enrichment of oxidative stress response genes in the neonatal gut, with six of nine reactive oxygen species (ROS)-scavenging enzyme mutants displaying neonatal-specific fitness defects **(Fig. S3a)**, highligting the importance of aerotolerance^21,24^. Furthermore, glyoxylate/dicarboxylate and propanoate metabolism emerged as enriched pathways (despite the absence of a canonical glyoxylate shunt^*25*^), suggesting roles in NADH generation through TCA cycle and redox balancing under transient oxygen exposure **(Fig. S3b)**^26,27^. Collectively, these findings reveal that *B. fragilis* integrates aerobic respiration, robust stress defenses, and complementary metabolic routes to withstand oxygen exposure in the neonatal gut, demonstrating a multifaceted strategy for early-life colonization success.

### Alpha-galactosylceramide is required for *B. fragilis* colonization in the infant gut

To pinpoint *B. fragilis*-specific factors underlying its distinct neonatal colonization pattern, we interrogated Tn-seq hits for genes unique to this species among ten *Bacteroidetes* genomes used in the colonization experiment **(Figs. 1a and 1e, and Supplementary Table 3)**. Of 575 species-specific genes (∼12.1% of the *B. fragilis* genome), 98 were required for colonization in neonates.

In light of the transiently microaerobic environment of the neonatal gut, we searched for colonization genes specifically induced upon oxygen exposure. Cross-referencing our Tn-seq screen with previously published transcriptome data **(Supplementary Table 1)**^26^ revealed only 3 out of 98 neonatal-specific fitness genes that were upregulated in response to oxygen, notably including *agcT* (BF638R_3264), encoding ceramide alpha-galactosyltransferase^28,29^. This enzyme synthesizes alpha-galactosylceramide (hereafter referred to as BfaGC, for *B. fragilis*-derived alpha-galactosylceramide), a well-characterized immunomodulatory sphingolipid known to regulate NKT cell homeostasis in large intestine^30-32^. Although the role of BfaGC in host immune regulation is well established, its functional relevance to *B. fragilis* physiology remains elusive. Sphingolipids are major membrane lipid components of *Bacteroidetes*, essential for maintaining membrane integrity and facilitating bacterial survival in the gut under environmental stress^33^. Furthermore, bacterial sphingolipids play pivotal roles in host-microbe interactions, influencing intestinal inflammation^34^ and host lipid metabolism^35,36^. Notably, core ceramide synthetic genes (*spt, cerR*, and *cerS*)^37^ are necessary for colonization in the adult gut **(Fig. 1f)**, whereas *agcT* is uniquely essential in neonates, suggesting that galactosylation of ceramide to form aGC provides a context-specific benefit distinct from the structural role of its precursor.

### BfaGC is a necessary and sufficient fitness factor of early gut colonization

To test the functional role of BfaGC, we introduced a 1:1 mixture of wild-type and Δ*agcT B. fragilis* into germ-free mice **(Fig. 2a)**. Deletion of *agcT* abolished BfaGC synthesis without affecting bacterial growth or upstream ceramide production **(Figs. S4a and S4b)**. While both strains persisted equally in adult mice, Δ*agcT* exhibited a sevenfold colonization defect in neonates following natural vertical transmission, consistent with Tn-seq results. This defect was not observed when colonization was initiated postnatally at day 7, indicating BfaGC is specifically required for vertical transmission and early establishment.

**Figure 2.**
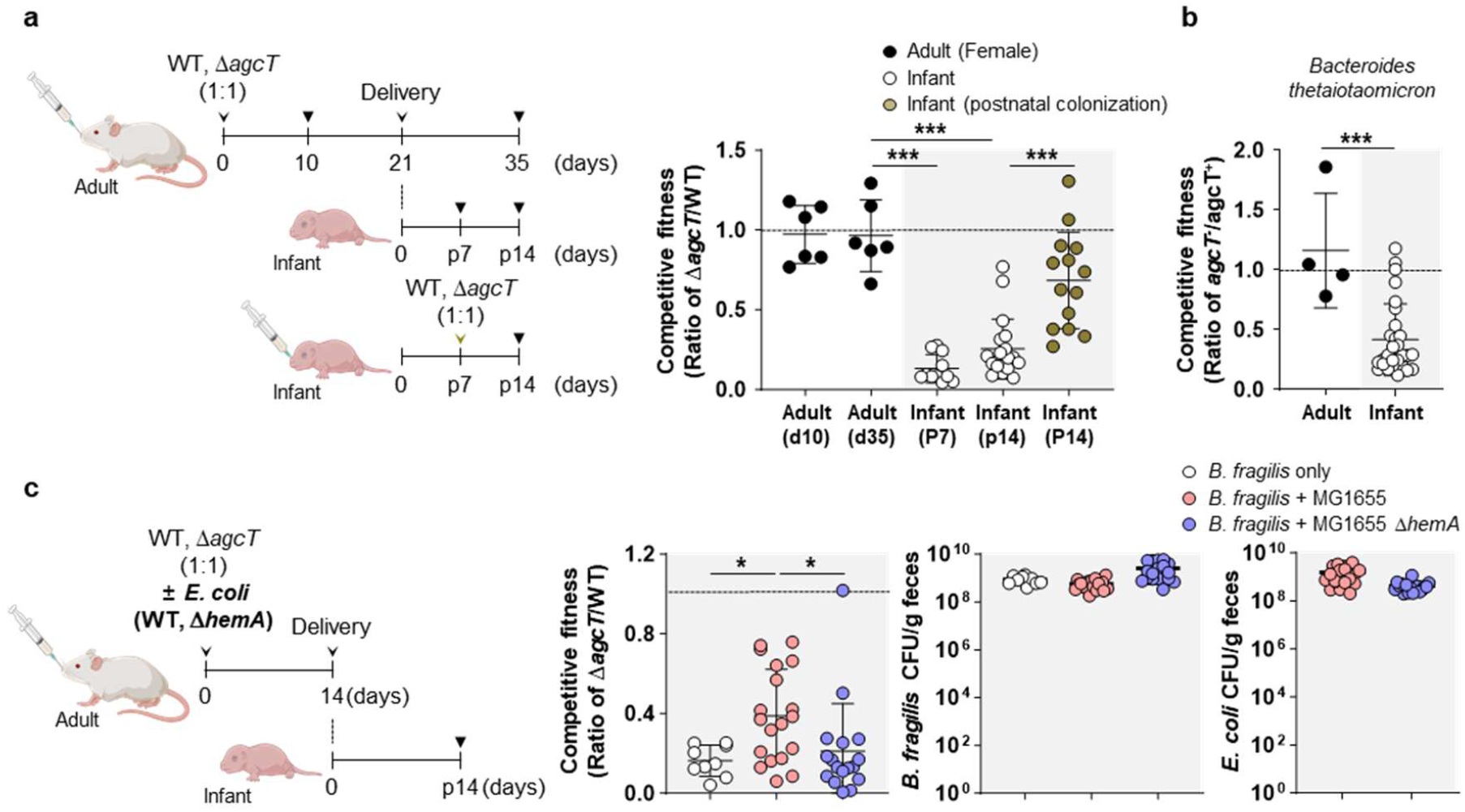
BfaGC enhances *B. fragilis* colonization in the neonatal gut by promoting aerotolerance. **a**, Germ-free Swiss-Webster mice were orally gavaged with a 1:1 mixture of wild-type and Δ*agcT* mutant *B. fragilis*, and bacterial abundances were measured in neonatal and adult mice. For postnatal colonization, infant mice were gavaged with the mixture at postnatal day 7 (p7). **b**, Competitive colonization assay of *Bacteroides thetaiotaomicron* strains lacking (*agcT*^-^), or overexpressing *agcT* (*agcT*^+^). Competitive fitness of *agcT* was calculated by denominating the abundance of *agcT*^-^ strain by that of *agcT* ^+^ strain. **c**, Germ-free pregnant mice were co-colonized with a 1:1 mixture of wildtype *B. fragilis* and its Δ*agcT* mutant, in the presence or absence of either wildtype *E. coli* or an *E. coli* Δ*hemA* mutant. The abundance of *B. fragilis* strains as well as *E. coli* stains were determined in the neonatal gut at p14. Shown are the means and SD.

To further test whether BfaGC alone is sufficient to promote early colonization, we engineered a non-aGC-producing species, *Bacteroides thetaiotaomicron*, to express *agcT*. Heterologous expression conferred aGC synthesis **(Figs. S4c and S4d)**. In competitive colonization assays, the *agcT*-expressing strain outcompeted its parental counterpart in neonatal mice, demonstrating that BfaGC production is sufficient to enhance early-life colonization **(Fig. 2b, Fig. S4e)**.

### BfaGC enhances the aerotolerance of *B. fragilis*

Previous transcriptomic analyses showed that *agcT* is upregulated in response to oxygen exposure^26^. Given the enrichment of oxidative stress response pathways during neonatal colonization and the established role of sphingolipids in such stress adaptation^33^, we hypothesized that BfaGC enhances *B. fragilis* aerotolerance. As expected, both *agcT* expression and BfaGC production increased following short-term oxygen exposure *in vitro* **(Fig. S5a)**. While wild-type and Δ*agcT* strains exhibited comparable growth under anaerobic and microaerobic conditions, Δ*agcT* strains displayed markedly reduced viability following aerobic challenge, which was fully restored by *agcT* complementation **(Figs. S6a and S6b)**. These findings indicate that BfaGC contributes to oxygen tolerance, likely supporting bacterial fitness under early-life oxygen stress.

To test whether BfaGC promotes early gut colonization by enhancing oxygen persistence *in vivo*, we performed co-colonization assays with *Escherichia coli*, a facultative anaerobe that consumes oxygen in the gut **(Fig. 2c)**. The presence of *E. coli* attenuated the fitness difference between wild-type and Δ*agcT* strains while maintaining the total *B. fragilis* level. This attenuation effect was abolished when co-colonized with *E. coli* Δ*hemA*, a strain deficient in heme synthesis and consequently incapable of aerobic respiration^38^, indicating that BfaGC-mediated colonization is contingent upon environmental oxygen levels. Together, these results suggest that BfaGC promotes *B. fragilis* persistence in the neonatal gut by enhancing tolerance to transient oxygen exposure.

### BfaGC stabilizes membrane integrity to support aerobic respiration

The neonatal gut is a metabolically dynamic niche marked by transient oxygen exposure and a predominance of facultative anaerobes. Given that ceramide is a major membrane lipid in *Bacteroidetes*, we hypothesized that its glycosylation into BfaGC alters membrane properties to support colonization in this partially oxygenated milieu. Because aerobic respiration is primarily mediated by membrane-bound complexes, we asked whether BfaGC contributes to respiratory adaptation and fitness.

Oxygen consumption assays revealed that the Δ*agcT* mutant exhibited decreased aerobic respiration compared to wild-type *B. fragilis*, while the Δ*cyd* strain, lacking cytochrome *bd* oxidase, showed no oxygen consumption at all **(Fig. 3a, Fig. S7a)**. The Δ*agcT* mutant exhibited a higher ATP/ADP ratio and elevated short-chain fatty acid (SCFA) production upon brief oxygen exposure, suggesting a sustained reliance on fermentative metabolism despite oxygen availability. **(Fig. S7b and 7c)**. This overflow metabolism of the Δ*agcT* mutant point to insufficient coupling of metabolic flux into the electron transport chain (ETC). Tn-seq analysis indicated a reconfiguration of respiratory chains in *B. fragilis*, with respiration shifting from sodium-coupled complexes (Nqr, V-type) in adults to proton-dependent systems (Nuo, F-type ATPase) in neonates **(Fig. S8)**^39^. This bioenergetic shift is further supported by transcriptomic data showing repression of Nqr operon upon oxygen exposure with concurrent induction of cytochrome *bd*^26^. As BfaGC is oxygen-inducible, we hypothesized that it supports this shift by enhancing proton retention and PMF generation.

**Figure 3.**
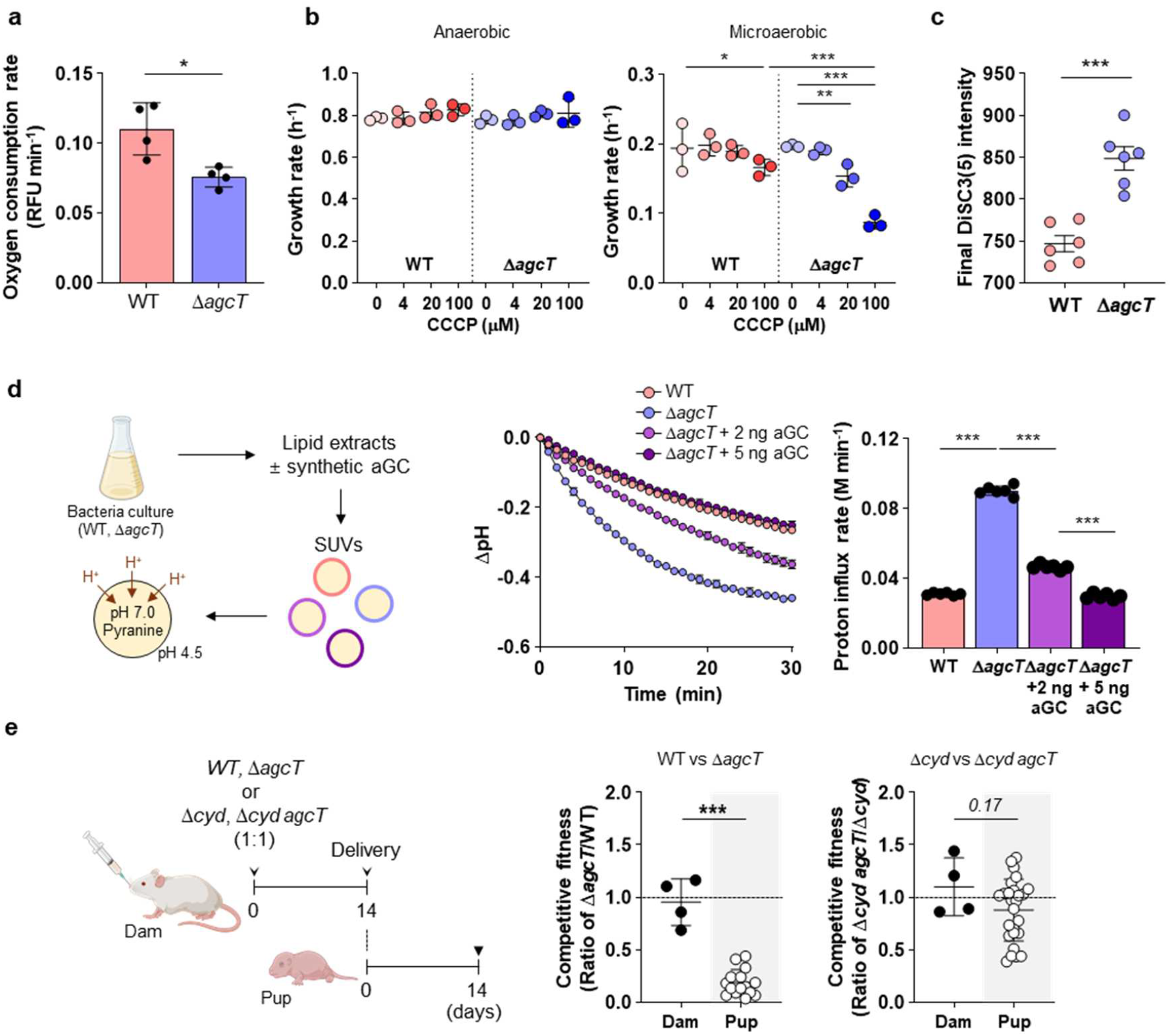
BfaGC enhances *B. fragilis* aerobic respiration by stabilizing membrane potential and proton retention. **a**, Relative oxygen consumption rate of WT and Δ*agcT* under microaerobic condition. **b**, Growth rates of WT and Δ*agcT* was measured in PYG medium in the presence of indicated amounts of CCCP under anaerobic and microaerobic condition. **c**, Membrane potential measurements using the DiSC3(5) fluorescence assay. Bacterial cells were treated with DiSC3(5) at 5 min and fluorescence was monitored for 40 min with excitation wavelength of 651 nm and emission wavelength of 675 nm. **d**, Proton permeability was measured using small unilamellar vesicles (SUVs) composed of lipid extracts from wildtype, Δ*agcT*, and Δ*agcT* supplemented with synthetic BfaGC (2 ng or 5 ng). SUVs were loaded with the pH-sensitive fluorescent dye pyranine (2 mM) and equilibrated in a neutral buffer (pH 7.0). To induce proton influx, SUVs were transferred to an acidic buffer (pH 4.5), and intravesicular pH changes were monitored over 30 min. Fluorescence was measured at excitation wavelengths of 410 nm and 454 nm, with emission at 510 nm. Intravesicular pH values were calculated using a standard calibration curve based on the fluorescence ratio (410/454 nm). The proton influx rates were determined by performing linear regression on time-resolved values obtained through inverse logarithmic transformation of measured pH. **e**, Competitive colonization assay of Δ*cyd* and Δ*cyd* agcT *B. fragilis* in neonatal mice. Germ-free mice were co-colonized with either a 1:1 mixture of WT and Δ*agcT* or a mixture of Δ*cyd* and Δ*cyd agcT*. Competitive fitness of the *agcT* gene was determined by calculating the ratio of Δ*agcT* to WT (Δ*agcT*/WT) or Δ*cyd agcT* to Δ*cyd* (Δ*cyd agcT*/Δ*cyd)* Shown are the means and SD.

To determine whether BfaGC contributes to proton motive force (PMF)-driven respiration, we exposed wild-type and Δ*agcT* strains to the protonophore CCCP. While both strains were unaffected under anaerobic conditions, Δ*agcT* exhibited heightened CCCP sensitivity under microaerobic conditions **(Fig. 3b)**. Consistently, membrane potential was significantly reduced in the Δ*agcT* mutant compared to wild-type **(Fig. 3c)**, indicating BfaGC as a key factor in sustaining PMF maintenance. Of note, PMF is typically generated by electron transport chain complexes, the activity of which remained unaffected by BfaGC (**Fig. S7d and 7e)**.

To sustain an efficient PMF generation, membrane lipids must exhibit exceptional proton impermeability, minimizing proton leakage via “proton hopping” through hydrogen-bonded networks and thus energy dissipation^40,41^. Glycolipids such as digalactosyldiacylglycerol were shown to reduce membrane proton permeability^42^. Building on this, we hypothesized that BfaGC similarly enhances membrane impermeability to stabilize PMF and support aerobic respiration under microaerobic conditions. To assess whether BfaGC modulates membrane proton permeability, we generated small unilamellar vesicles (SUVs) from lipid extracts of wild-type and Δ*agcT B. fragilis* cultures **(Fig. 3d)**. When exposed to an acidic buffer, Δ*agcT*-derived SUVs showed significantly increased proton influx compared to the wild-type counterpart. Incorporation of synthetic BfaGC into Δ*agcT*-derived liposomes restored proton impermeability to wild-type levels, indicating that BfaGC directly enhances membrane proton retention. This role of BfaGC in enhancing membrane proton impermeability was further validated using synthetic lipid vesicles **(Fig. S9)**, demonstrating that physiological aGC/ceramide ratios are sufficient to modulate proton influx rates.

To determine whether the colonization advantage conferred by BfaGC is dependent on aerobic respiration, we assessed the fitness of Δ*cyd* and Δ*cyd agcT* mutants, both lacking cytochrome *bd* oxidase, under microaerobic conditions. Both strains exhibited comparable viability **(Fig. S6c)**, suggesting that the contribution of BfaGC to aerotolerance requires functional respiration. Consistently, in neonatal colonization assays, the Δ*agcT* mutant displayed a significant fitness defect relative to wild-type, whereas no difference was observed between Δ*cyd* and Δ*cyd agcT* strains **(Fig. 3e)**. These findings demonstrate that the beneficial effect of BfaGC on neonatal colonization is contingent upon the respiratory adaptation.

### BfaGC is required for temporal niche acquisition of *B. fragilis*

Sphingolipids are widely produced among *Bacteroidetes*, yet only a few species, including *B. fragilis*, synthesize aGC^43^. In contrast, the majority of *Bacteroidetes* species generate a different class of glycosphingolipid, phosphoinositol dihydroceramide (PI-cer) **(Fig. 4a)**^44^. This biochemical divergence among gut microbiota raises key evolutionary questions regarding the distinct function and selective advantages conferred by these lipids.

**Figure 4.**
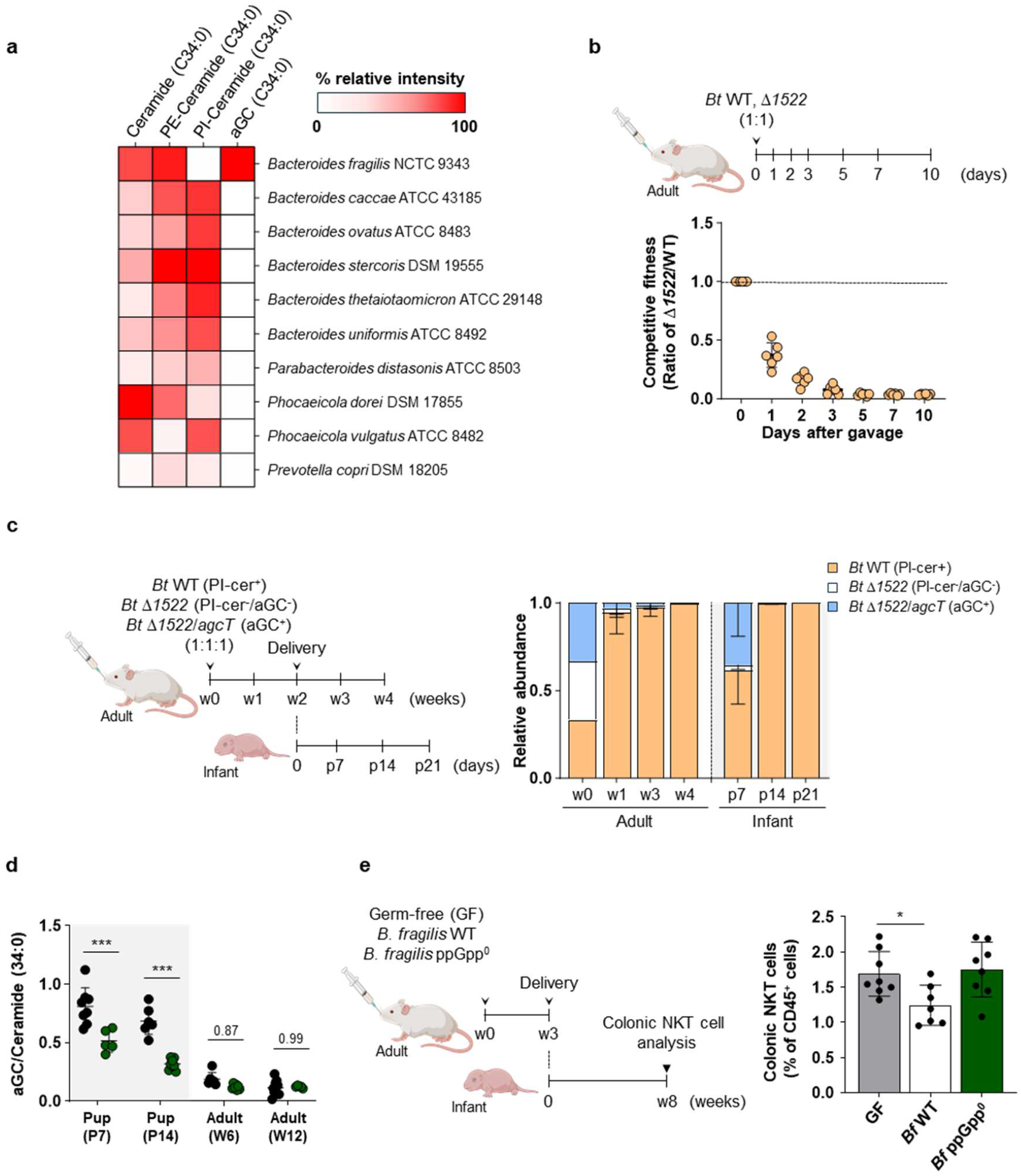
Oxygen-responsive BfaGC mediates temporal niche acquisition and immune modulation by *B. fragilis* in the neonatal gut. **a**, The synthesis of ceramide derivatives across different *Bacteroidetes* species. The synthesis of alpha-galactosylceramide (aGC) is restricted to *B. fragilis*, while other species predominantly synthesize phosphoinositol-ceramide (PI-cer). **b**, Competition colonization assay between wildtype *B. thetaiotaomicron* and PI-cer-deficient mutant (Δ*1522*) strains in adult mice. Ther abundance was monitored for 10 days after co-colonization by measuring colony-forming units (CFUs). **c**, Tripartite competition assay comparing strains producing PI-ceramide (PI-cer^+^), aGC (aGC^+^), or neither (PI^-^, aGC^-^). Their abundance was measured by qRT-PCR using their specific-primers. **d**, Quantification of aGC levels in wild-type and ppGpp^0^ mutant *B. fragilis* strains under neonatal (at postnatal day 7 and 14) and adult gut conditions (at 6- and 12-weeks old) **e**, Flow cytometry analysis of colonic NKT cell frequency in germ-free (GF), wild-type *B. fragilis* (WT) at week 8. Shown are the means and SD.

To directly investigate the selective pressures governing aGC and PI-cer biosynthesis, we conducted co-colonization experiments comparing wild-type *B. thetaiotaomicron* and its PI-cer-deficient mutant (Δ*BT1522*) **(Fig. 4b)**. Δ*BT1522* exhibited a pronounced fitness defect in the adult gut, whereas *B. fragilis* Δ*agcT* did not exhibit such defect **(Fig. 2a)**, indicating that PI-cer is critical for colonization in the adult gut, whereas aGC is dispensable under these conditions. In contrast, Δ*agcT* displayed a significant fitness cost in the neonatal gut, demonstrating that aGC is crucial for early-life colonization. This divergence in fitness advantage underscores the distinct evolutionary pressures acting on these sphingolipids: The adult gut represents a stable, long-term selective landscape that favors the broad conservation of PI-cer across *Bacteroidetes*, whereas aGC-mediated adaptation appears tailored to a transient neonatal phase, where rapid microbial colonization precedes the onset of strict anaerobiosis. This temporal constraint likely contributes to the differential conservation of these biosynthetic pathways, with PI-cer optimizing persistence across the host lifespan and aGC facilitating successful niche acquisition in early life.

To further elucidate this distinction, we conducted a tripartite competition assay using *B. thetaiotaomicron* strains producing PI-cer (PI-cer^+^), aGC (aGC^+^), or neither (PI^-^, aGC^-^) **(Fig. 4c)**. In the adult gut, the PI-cer^+^ strain outcompeted both the aGC^+^ and PI^-^, aGC^-^ strains, constituting 96% of the population within one week and 99% by week 4. This finding confirms that PI-cer biosynthesis provides a substantial fitness advantage in the adult gut microbiota. Conversely, in the neonatal gut, the aGC^+^ strain accounted for 35.5% of the total population, far exceeding its representation in the adult gut. This result reinforces the notion that aGC production is a critical determinant for early-life microbial niche acquisition.

Beyond its temporal role, BfaGC also conferred a spatial fitness benefit in oxygenated regions of the adult gut. Lipidomic profiling and regional fitness mapping revealed that both BfaGC abundance and *agcT*-dependent fitness peaked in the ileum, an oxygen-rich compartment relative to the colon **(Fig. S10)**. Collectively, these findings establish BfaGC as a key determinant of *B. fragilis* persistence across distinct oxygen landscapes, conferring competitive advantages in both temporal and spatial niche acquisition, highlighting its broader significance in shaping microbial fitness within dynamic gut environments.

### Oxygen-induced BfaGC drives temporal immunomodulation by *B. fragilis*

In addition to its metabolic role in facilitating early colonization, BfaGC functions as a temporally restricted immunomodulatory lipid that links microbial adaptation to host immune development. Bf aGC was shown to regulate colonic NKT cell proliferation throughout life^30-32^. This effect is confined to the neonatal period, coinciding with both transient oxygen exposure and peak *B. fragilis* colonization^45^. Given this synchrony, we hypothesized that oxygen-triggered BfaGC production serves as a co-evolved mechanism aligning microbial niche acquisition with host immune imprinting.

To assess this, we used a stringent response-deficient *B. fragilis* mutant (ppGpp^0^, Δ*BF9343_0658, BF9343_2136*), which has impaired oxidative stress adaptation and fails to upregulate BfaGC in response to oxygen **(Fig. S5b)** ^46-48^. In monocolonized neonates, the ppGpp^0^ strain did not induce BfaGC production **(Fig. 4d)** despite establishing colonization levels comparable to those of the wild-type strain. In accordance with the loss of BfaGC induction, animals monocolonized with the ppGpp^0^ mutant failed to regulate colonic NKT cell levels, in contrast to the wild-type *B. fragilis* monocolonized group **(Fig. 4e)**. Notably, despite this immunological divergence, the ppGpp^0^ strain exhibited comparable CFUs and BfaGC levels in the adult gut to the wild-type strain, confirming that the immunomodulatory effects of BfaGC are specific to the neonatal period and not merely a consequence of bacterial load or persistence.

These findings demonstrate BfaGC as a species-specific sphingolipid that equips *B. fragilis* to exploits the neonatal gut through enhanced respiratory fitness and membrane stabilization. By coupling metabolic adaptation with temporally restricted immune modulation, BfaGC exemplify a co-evolved microbial strategy for early-life niche acquisition and long-term symbiosis.

## Discussion

To assure their coexistence with the host over generations, symbionts of mammalian GI tract must find their way to endure challenges caused by rapid physiological changes of the host. Our study reveals a previously unrecognized metabolic adaptation that enables *B. fragilis* to efficiently colonize the neonatal gut, an environment characterized by transient oxygen exposure and dynamic host development. While strict anaerobes are traditionally thought to rely solely on fermentation and anaerobic respiration, we demonstrate that *B. fragilis* engages in oxygen-dependent respiration during early colonization. This transition involves fundamental bioenergetic reconfiguration, wherein *B. fragilis* shifts from sodium motive force (SMF)-driven respiration in the adult gut to PMF-driven respiration in neonates. The ability to leverage cytochrome *bd* oxidase-dependent respiration in response to fluctuating oxygen levels provides *B. fragilis* with a distinct colonization advantage over obligate anaerobes that lack such respiratory flexibility.

A central finding of our study is the identification of BfaGC as a critical determinant of this bioenergetic transition. While bacterial sphingolipids have been primarily studied in the context of host physiological modulation^31,34-36^, our data reveal a distinct role for BfaGC in stabilizing membrane potential and minimizing proton leakage, thereby supporting PMF maintenance and aerobic respiratory efficiency. This function parallels, but is mechanistically distinct from, other known lipid-mediated respiratory adaptations, such as cardiolipin-driven electron transport optimization in both bacteria and mitochondria^49,50^. The biophysical role of BfaGC aligns with recent work in *Escherichia coli* linking membrane viscosity to respiration rate^51^, highlighting a broader principle wherein membrane composition dynamically tunes bacterial bioenergetics in response to environmental stress.

Beyond its metabolic function, BfaGC biosynthesis appears to represent an evolutionary strategy enabling *Bacteroides fragilis* to colonize transiently oxygenated niches during early life. In contrast, phosphoinositol dihydroceramide (PI-cer) supports persistence in the strictly anaerobic adult gut^44^. Given that both lipids derive from a common ceramide scaffold and are exclusively distributed among gut *Bacteroidetes*—with species producing BfaGC lacking PI-cer biosynthesis and *vice versa*—the persistence of the aGC trait across generations, despite its limited utility in adult environments, presents an evolutionary paradox. A plausible resolution lies in the spatial ecology of *B. fragilis*, which preferentially localizes to partially oxygenated mucosal regions such as the ileum, epithelium, and even lamina propria^52-54^, where aGC-mediated aerotolerance may confer a sustained fitness advantage beyond the neonatal window. This positional specialization may not only facilitate long-term maintenance of BfaGC-expressing lineages, but may also have contributed to the emergence of enterotoxigenic *B. fragilis* (ETBF), a variant adapted to exploit oxygen-exposed mucosal environments^54,55^. These findings highlight how a single lipid-mediated adaptation can integrate temporal niche acquisition, spatial persistence, and evolutionary diversification within the gut microbiota.

Co-adaptive microbial traits that promote colonization while modulating host physiology are widely observed in host–microbe interactions. *B. fragilis* polysaccharide A (PSA) supports bacterial persistence and induces regulatory T cell differentiation^19,56^; commensal bile salt hydrolases deconjugate host bile acids, mitigating their toxicity while influencing host lipid metabolism^57,58^; and structural variants of commensal lipopolysaccharide attenuate innate activation and enhance resistance to antimicrobial peptides^59,60^. Within this broader paradigm, BfaGC represents a temporally restricted instance in which both microbial and immunological functions are synchronously aligned during early life^31^. Its coordinated role during early life offers concrete evidence of microbial–host co-evolution, illustrating how specific microbial traits can be temporally calibrated to coincide with key developmental stages in the host, thereby reinforcing the importance of timing in shaping stable and beneficial host–microbe relationships.

Together, our findings reveal BfaGC as a metabolically and evolutionarily significant glycolipid that enables *B. fragilis* to navigate the unique physiological landscape of the neonatal gut. By linking membrane biophysics to respiratory efficiency and coupling microbial colonization with immune modulation, BfaGC exemplifies how bacterial metabolites can be temporally tuned to align with critical host developmental windows. This work expands the current understanding of microbial adaptation by demonstrating that even single-molecule traits can mediate complex co-evolutionary outcomes.

## Methods

### Mouse

All animal procedures were conducted under the supervision of the Harvard Center for Comparative Medicine and maintained by the Institutional Animal Care and Use Facility. Mice used in the experiments were 1–12 weeks old (including neonates), with all experimental groups being age-matched. The mice were housed in a controlled environment with a 12-hour light/dark cycle, a temperature of 21°C, and 50% humidity.

Germ-free mice were bred and maintained within inflatable plastic isolators. Gnotobiotic mice (either monocolonized or co-colonized for competition assay) were generated by orally gavaging breeding pairs with the bacterial strains, after which they were kept in isolators to produce offspring (F1 and subsequent generations) for experimental use. Stool samples from both GF and monocolonized mice in the isolators were regularly cultured on plates under both aerobic and anaerobic conditions to ensure sterility and the absence of contamination.

### Bacteria culture

For bacterial cultures, frozen stocks maintained at -80°C were streaked onto Brain Heart Infusion (BHI) agar plates. A single colony was inoculated into deoxygenated peptone-yeast extract-glucose (PYG) broth (2% proteose peptone, 0.5% yeast extract, 0.5% NaCl, supplemented with 0.5% glucose, 0.5% K_2_HPO_4_, 0.05% l-cysteine, 5 mg L^-1^ hemin, and 2.5 mg L^-1^ vitamin K1) within an anaerobic chamber. For further bacterial growth, the overnight cultures were inoculated into media under the specified conditions. For metabolomics analysis, bacteria were grown in minimal medium (0.1% (NH_4_)_2_SO_4_, 0.1% Na_2_CO_3_, 0.09% KH_2_PO_4_, 0.09% NaCl, 26.5 mg L^-1^ CaCl_2_·2H_2_O, 20 mg L^-1^ MgCl_2_·6H_2_O, 10 mg L^-1^ MnCl_2_·4H_2_O, 1 mg L^-1^ CoCl_2_·6H_2_O, 0.05% l-cysteine, 5 mg L^-1^ hemin, 2.5 mg L^-1^ vitamin K1, 2 mg L^-1^ FeSO_4_·7H_2_O, and 5 μg L^-1^ vitamin B12), with the addition of the indicated carbon sources.

Antibiotics for selection were used in the following concentrations: 100 µg ml^-1^ ampicillin for *Escherichia coli*, and 10 µg ml^-1^ erythromycin, 200 µg ml^-1^ gentamycin, and 100 ng ml^-1^ or the indicated amounts of anhydrotetracycline for *Bacteroides fragilis* and *Bacteroides thetaiotaomicron*.

For colonization experiments using 10-member *Bacteroidetes* community, the following strains were used: *B. fragilis* NCTC9343, *Bacteroides caccae* ATCC 43185, *Bacteroides thetaiotaomicron* VPI-5482, *Bacteroides uniformis* ATCC 8492, *Bacteroides stercoris* DSM 19555, *Phocaeicola vulgatus* ATCC 8482, *Phocaeicola dorei* DSM 17855, *Bacteroides ovatus* ATCC 8483, *Prevotella copri* DSM 18205, and *Parabacteroides distasonis* ATCC 8503.

### Construction of *Bacteroides* knockout/overexpression mutants

Primers used for this study were described in Supplementary table 4. Δ*agcT* (Δ*BF9343_3149*) and ppGpp^0^ (Δ*BF9343_0658, BF9343_2136*) mutant *B. fragilis* and Δ*BT_1522 B. thetaiotaomicron* were constructed using the pLGB13 suicide vector^61^. For overexpression, the *agcT* gene or *relA* gene (BF9343_0658) was cloned into the pNBU2_erm-TetR-P1T_DP-GH023 vector under either their native promoter or the TetR promoter (anhydrotetracycline-inducible)^62^.

### Construction of transposon mutant library

Transposon mutagenesis was performed via conjugation between *Bacteroides fragilis* 638R recipient cells and *Escherichia coli* S17-1 λpir donor cells harboring the transposon-containing plasmid pSAM-Bt^63^. For this process, 2 mL of overnight-cultured *B. fragilis* was inoculated into 800 mL of fresh PYG medium, while the donor *E. coli* was grown in 50 mL of LB medium supplemented with ampicillin. The recipient and donor cells were cultured until reaching exponential growth phases (*B. fragilis* OD600 0.05–0.1, *E. coli* OD600 ∼0.5), and subsequently mixed at a ratio of approximately 10:1 based on optical density. The cell suspension was harvested by centrifugation at 8,000 × *g* for 5 minutes and resuspended in ∼5 mL of fresh PYG medium. The resulting cell mixture was spotted onto BHI agar plates and incubated aerobically for 16 hours. Post-incubation, the spots were scraped off the plates, resuspended in 5 mL of PYG medium, and spread onto BHI agar plates containing erythromycin and gentamicin to counter-select for *B. fragilis* transconjugants harboring the transposon. The plates were incubated under anaerobic conditions for 2–3 days. To estimate the total size of the mutant library, aliquots of the conjugated cell suspension were serially diluted, plated on selective agar, and colony-forming units (CFUs) were enumerated. The total library size was determined based on CFU calculations. The resultant transconjugants were resuspended in PYG medium containing 15% glycerol and immediately stored at −80°C. For subsequent experiments, individual aliquots were thawed and cultured as needed.

### *In vivo* selection of transposon mutant library

The transposon mutant library was cultured anaerobically in fresh PYG medium for 2 hours, corresponding to approximately 2–3 replication cycles, to minimize potential growth rate biases among mutants. The library was grown to an optical density (OD600) of 1.0. Aliquots of 200 μL, containing approximately 2 × 10^8^ mutant cells, were prepared for oral gavage. Pregnant germ-free Swiss-Webster female mice were administered the mutant library via oral gavage at embryonic days 7–9 (E7–9). Fecal samples were collected from the adult female mice on the day of delivery, as well as from their offspring on postnatal day 10.

### Transposon sequencing using INseq method

Following selection, genomic DNA was isolated from the mutant population. The extracted gDNA was digested with the restriction enzyme MmeI, which specifically cleaves both termini of the transposon cassette. The digestion reaction was carried out at 37°C for 2 hours and subsequently heat-inactivated at 65°C for 20 minutes. The digested DNA was resolved via agarose gel electrophoresis, and a fragment approximately 1.4 kb in size—corresponding to the transposon cassette—was excised based on a molecular size marker. DNA was purified from the gel using the Monarch Gel Extraction Kit, with final elution in nuclease-free water. Adaptor ligation was performed by incubating the purified DNA with a 0.5 μM adaptor containing sample-specific barcode, and T4 DNA ligase at 20°C for 2 hours. The ligated DNA fragments were PCR-amplified using Lib_5 and Lib_3 primers. The PCR protocol included an initial denaturation step at 98°C for 30 seconds, followed by as few amplification cycles as necessary to remain within the linear range of amplification (98°C for 10 seconds, 63°C for 10 seconds, and 72°C for 10 seconds; 20 cycles), with a final hold at 4°C. The amplified PCR products were purified from agarose gels, and their concentration and integrity were assessed using the Agilent TapeStation High Sensitivity D5000 assay. Sequencing libraries were prepared and analyzed using the Illumina NextSeq 500 platform.

### Data Processing and Statistical Analysis

Quality control, adapter sequence trimming, and removal of transposon sequences from the reads were performed using Trimmomatic^64^. Processed reads were aligned to the *B. fragilis* 638R reference genome using Bowtie2 v2.4.2 with default parameters^65^. Mapped reads were quantified by FeatureCounts^66^, assigning read counts to corresponding open reading frames (ORFs). Differential abundance analysis of read counts was conducted using either DESeq2^67^ or zero-inflated negative binomial (ZINB) models, depending on the experimental group. An enrichment analysis of functional terms for genes of interest was carried out using KEGG gene set enrichment analysis^68^.

To identify essential genes for in vitro growth, insertion sites from sequencing data were mapped to annotated gene regions. Insertion events were categorized by matching their positions to the start and end coordinates of ORFs. Unmatched insertion sites were excluded from further analysis. Read counts for each gene were aggregated and analyzed using a Bayesian modeling approach^69^. The posterior probabilities for transposon insertion at each gene were estimated using PyMC3. For each gene, insertion probability was modeled using a uniform Beta(1,1) prior, and observed read counts were assumed to follow a Binomial likelihood. Posterior inference was performed using the No-U-Turn Sampler (NUTS) with default PyMC3 settings. All MCMC chains demonstrated satisfactory convergence (≈ 1.0). Gene-level posterior probabilities were then log-transformed, yielding a distinctly bimodal distribution **(Fig. S1b)**, with the lower peak corresponding to essential genes (low probabilities of transposon insertions) and the upper peak representing non-essential genes. To objectively separate these two groups, a two-component Gaussian mixture model (GMM) was fitted to the distribution, and the cutoff was determined at the intersection point of the two Gaussian components, where the posterior probabilities of belonging to either category are equal. To further enhance stringency, only genes that additionally exhibited a high component membership probability (P > 0.95) in the essential cluster were ultimately classified as essential.

For identifying genes essential for colonization in the adult gut, DESeq2 was employed to analyze differential abundance of read counts between samples collected from the initial mutant library and those recovered from adult mouse fecal samples. Read count normalization was performed by calculating size factors based on the total reads per sample. Results were adjusted for multiple testing using the Benjamini-Hochberg method, and genes with an adjusted *p*-value <0.05 were considered significantly enriched.

In the neonatal gut, ZINB models were used to account for excess zeros and overdispersion in read count data. Samples from adult and neonatal mice were compared using a generalized linear mixed-effects model (glmmTMB package). Gene-wise estimates and p-values for group-specific differences were extracted from the model. False discovery rates (FDR) were calculated to identify genes with significant changes in fitness, using an FDR cutoff of 0.05. Functional enrichment analysis was then performed on significant gene sets to identify critical pathways contributing to bacterial survival in the neonatal environment.

### Competitive colonization assay

The colonization fitness of different *B. fragilis* or *B. thetaiotaomicron* strains was assessed by a competitive colonization assay. Erythromycin-sensitive and erythromycin-resistant strains (engineered via pNBU2 cassette insertion) were cultured in liquid PYG medium until they reached exponential growth phase (OD600 ∼1.0). Equal volumes of each strain, adjusted to identical optical densities, were mixed prior to oral gavage into germ-free Swiss Webster mice. A fraction of the input mixture (used for gavage) and stool samples collected at defined time points were plated on BHI agar with and without erythromycin to distinguish between the two strains. Colony-forming units (CFUs) were enumerated, and the relative proportions of the two strains in each sample were calculated. The normalized ratio of the strains in stool samples was determined by comparing their proportions to those of the initial inoculum.

For tripartite competition assays, the proportions of individual strains were determined using quantitative real-time PCR (qRT-PCR). Stool samples were collected at defined time points, including the initial inoculum, and genomic DNA (gDNA) was extracted. Strain-specific primers were employed to quantify the abundance of each strain, enabling accurate calculation of their relative proportions within the microbial community.

### RNA extraction and quantitative real-time reverse transcription PCR (qRT-PCR)

RNA was extracted from the bacterial cells cultured under the indicated condition using the Direct-zol RNA Miniprep kits (Zymo research) following the manufacturer’s instructions. Total RNA (1,000 ng) from each sample was used to synthesize cDNA using the qScript cDNA Synthesis Kit (Quantabio). Twenty-fold diluted cDNA was subjected to real-time PCR amplification using the iTaq Universal SYBR Green One-step Kit (BioRad) with specific primers in a CFX96™ Real-Time System (BioRad, Hercules, CA, USA).

### Colonic NKT cell analysis

Germ-free (GF) mice and mice monocolonized from birth with either WT *B. fragilis* or its isogenic ppGpp^0^ mutant were euthanized for analysis. The large intestines were excised, with surrounding adipose tissue carefully removed. The intestines were opened longitudinally, thoroughly cleared of fecal content, sectioned into 1-inch segments, and subjected to two washes in HBSS containing 2 mM EDTA for 30 minutes at 37°C. Following epithelial cell removal, the tissues were rinsed briefly in HBSS, then incubated in RPMI 1640 containing 10% FBS, collagenase type VIII (1 mg ml^-1^), and DNase I (0.1 mg ml^-1^) for 45 minutes at 37°C with continuous agitation. The resultant digested tissues were resuspended in FACS buffer (PBS with 2% FBS and 1 mM EDTA), filtered sequentially through 70 μm and 40 μm strainers, and prepared for subsequent flow cytometry analysis.

For flow cytometry staining, the following reagents were employed: APC-labelled mCD1d tetramer (unloaded or PBS-57 loaded; 1:500; NIH Tetramer Core Facility), anti-mouse MHC class II-PE/Cy7 (1:1000), TCRβ–PE (1:300), CD45–PerCP–Cy5.5 (1:300; Biolegend), and a cell viability dye (Fixable Viability Dye eFluor 780, 1:1,000; ThermoFisher). Samples were stained for 30 minutes at 4°C, followed by washing with cold FACS buffer. Flow cytometry was conducted using a BD FACSFortessa system (BD Biosciences), with gating performed based on forward and side scatter, singlet population, and viability. The frequencies of TCRβ_+_CD1d tetramer^+^ cells within the gated total CD45^+^ population were quantified as the target population. Data analysis and quantification were performed using FlowJo V10 software (BD Biosciences).

### Lipidomics and Metabolomics analyses

#### Sample Preparation

For lipidome analysis, bacterial cell pellets (harvested from 1 ml of OD 1.0 cultures) or mouse fecal and colonic content samples (∼20 mg) were extracted using an one phase extraction with 100% MeOH (for bacterial cells) and the Matyash method (for mouse samples; MeOH:MTBE:Water = 3:10:2.5). Perdeuterated (d_35_) β-galactosylceramide (Matreya) was included as an internal standard for normalization.

For cellular metabolite analysis, bacterial cultures grown in 50 ml of minimal media supplemented with 0.5% glucose were harvested, and metabolites were extracted using cold extraction buffer (MeOH:Acetonitrile:Water = 40:40:20 with 0.1% formic acid) at -20°C overnight. The extracts were centrifuged at 10,000 × *g* for 10 min, dried using a SpeedVac at 4°C, and reconstituted in 60 μl of MeOH for analysis.For short-chain fatty acid (SCFA) analysis, the spent media were subjected to chemical derivatization using 3-nitrophenylhydrazine (3-NPH). Briefly, 30 μl of medium was mixed with 30 μl of 5 μg ml^-1^ internal standard (d_4_-acetic acid, d_5_-propionic acid, d_7_-butyric acid in 100% acetonitrile), followed by the sequential addition of 20 μl of 200 mM 3-NPH in 50% acetonitrile and 20 μl of a mixture containing 120 mM *N*-(3-dimethylaminopropyl)-N′-ethylcarbodiimide hydrochloride and 6% pyridine in 50% acetonitrile. After incubation at 40°C overnight, the reaction was chilled on ice, diluted with 100 μl of water, and subsequently diluted 10-fold in 25% acetonitrile prior to LC-MS analysis.

### UHPLC-MS and MS/MS Conditions

Mass spectrometry analyses were performed using an ultra-high-performance liquid chromatography-tandem mass spectrometry (UHPLC–MS/MS) system (Thermo Scientific Vanquish RP-UPLC coupled with a Q Exactive Orbitrap). The analysis was conducted in both negative and positive ion modes with the following parameters: Spray voltage: 3.25 kV; Sheath gas: 40 AU; Auxiliary gas: 8 AU; Capillary temperature: 350°C; Auxiliary gas heater temperature: 350°C; S-lens RF level: 65.0 AU; Mean collision energy: 22.5 AU.

For lipidomics analysis, chromatographic separation was performed using a YMC-Triart C8 column (2.1 mm × 100 mm × 3 μm) under the following gradient elution conditions: **(0-2 min)** 50% 2-propanol, 10% acetonitrile, 2.5% formic acid, **(2-5 min)** Linear increase to 85% 2-propanol, 10% acetonitrile, 2.5% formic acid, **(5-12 min)** Isocratic hold, **(12-20 min)** Immediate re-equilibration to the initial conditions and hold. The column was maintained at 40°C, with a flow rate of 200 μL min^-1^.High-resolution mass spectrometry (MS1 scan resolution = 70,000 at m/z 200) was used in combination with top 10 data-dependent acquisition (DDA) at R = 17,500, using an isolation window of 1.0 m/z. Alternatively, a parallel reaction monitoring (PRM) approach was employed to selectively target ceramides and their derivatives, including aGC and PI-cer.

For cellular metabolite analysis, chromatographic separation was performed using an Acquity UPLC BEH Amide XP column (2.1 mm × 150 mm, 1.7 μm) under the following gradient conditions: **(0-2 min)** 95% acetonitrile with 10 mM ammonium acetate and 10 mM ammonium hydroxide (isocratic), **(2-15 min)** linear decrease to 20% acetonitrile with the same buffer composition, **(15-16 min)** isocratic hold, **(16-20 min)** return to initial conditions, followed by **(20-30 min)** re-equilibration. High-resolution mass spectrometry (MS1 scan resolution = 120,000 at m/z 200) was performed over an m/z range of 70-1,000.

For SCFA analysis, chromatographic separation was performed using an Agilent Zorbax C18 column (4.6 mm × 75 mm, 1.8 μm) at 40°C with a flow rate of 0.6 mL min^-1^. The gradient elution conditions were as follows: **(0-3 min)** 25% acetonitrile with 0.05% formic acid (isocratic), **(3-8 min)** linear increase to 97.5% acetonitrile with 0.05% formic acid, **(8-14 min)** isocratic hold, and **(14-20 min)** re-equilibration to initial conditions. High-resolution mass spectrometry (MS1 scan resolution = 70,000 at m/z 200) was performed over an m/z range of 100–700, and targeted detection of 3-NPH-derivatized SCFAs and internal standards was achieved using parallel reaction monitoring (PRM) at R = 17,500.

#### Data analysis

Data acquisition was performed using Xcalibur 4.0 (Thermo Fisher Scientific). Relative quantification for all LC-MS-based analyses—including lipidomics, cellular metabolomics, and SCFA profiling—was carried out by integrating MS1 peak areas. For lipidomic and metabolomic datasets, values were normalized to the recovery of internal standards and either sample mass (for fecal and intestinal contents) or optical density (OD) of bacterial cultures. For SCFA measurements, quantification was normalized to the volume of collected culture supernatant, using stable isotope-labeled standards for calibration.

### Determination of oxygen consumption rates of *B. fragilis*

Oxygen consumption rates (OCRs) of wild-type, Δ*agcT*, and Δ*cyd* mutant *B. fragilis* cells cultured under microaerobic conditions were measured using the Extracellular Oxygen Consumption Assay (ab197243, Abcam), following the manufacturer’s instructions with minor modifications. Bacterial cultures were harvested, immediately resuspended in minimal medium supplemented with 0.5% glucose, and diluted to an OD_600_ of 1.0. Two hundred microliters of the cell suspension were aliquoted into black-walled 96-well plates, followed by the addition of 15 μL of the extracellular O_2_ consumption reagent. Two drops (∼100 μL) of pre-warmed high-sensitivity mineral oil were added to each well to prevent oxygen diffusion. Fluorescence was monitored every 2 minutes for 2 hours using an excitation wavelength of 380 nm and an emission wavelength of 650 nm. Cells not treated with the dye were used as a negative control, and fluorescence values were normalized by subtracting the signal of dye-free samples. Cell-free medium (either aerobically incubated or deoxygenated) was used as an additional control to gauge baseline oxygen levels. Oxygen consumption rate (OCR) was determined by performing linear regression on the fluorescence intensity values over time. The slope of the linear portion of the time vs fluorescence plot (expressed as RFU min^-1^) was used as a measure of OCR for each replicate well.

### Liposome Preparation and Proton Permeability Measurement of Liposomes

Wild-type *B. fragilis* and its isogenic Δ*agcT* mutant were cultured under microaerobic conditions until reaching an optical density of ∼0.5. Lipids were extracted from 10 mL of bacterial culture using the Matyash method (MeOH:MTBE:Water=3:10:2.5). The extracted lipids, with or without supplementation of synthetic BfaGC (17:0/17:0) (SB2211)^30^ at the indicated concentrations, were evaporated to dryness under a nitrogen gas stream in a glass test tube.

In parallel, liposomes were also generated using defined synthetic lipid mixtures composed of 1,2-dioleoyl-sn-glycero-3-phosphatidylcholine (DOPC) (Cayman, 15098), C16 Dihydroceramide (d18:0/16:0) (Avanti Research, 860634), and C16 Galactosyl(α) Dihydroceramide (d18:0/16:0) (Avanti Research, 860730). Each lipid was prepared as a 2 mM stock solution, and 30 μL of DOPC and 20 μL of a ceramide/aGC mixture (in the indicated molar ratio) were combined and dried under nitrogen prior to rehydration and vesicle formation.

To generate liposome suspensions, the dried lipid film was then rehydrated with 300 μL of buffer solution (2 mM trisodium 8-hydroxypyrene-1,3,6-trisulfonate (pyranine) (ThermoFisher Scientific, L11252), 5 mM KH_2_PO_4_, 100 mM KCl, adjusted to pH 7.0 with KOH) at 50°C for 10 minutes. The rehydrated lipid mixture was sonicated for 20 minutes using an ultrasonic cleaner. To remove disordered liposomes, the suspension was centrifuged at 5000 × *g* for 10 minutes, followed by dialysis against pyranine-free buffer (5 mM KH_2_PO_4_, 100 mM KCl, adjusted to pH 7.0 with KOH) to eliminate any extraliposomal pyranine.

Proton permeability of liposomes was assessed as previously described with modifications. Each liposome suspension (50 μL) was mixed with 150 μL of acidic buffer (5 mM KH_2_PO_4_, 100 mM KCl, adjusted to pH 4.5 with HCl). Fluorescence emission at 510 nm was monitored over 30 minutes using a fluorescence spectrophotometer. The internal pH of the liposomes was determined based on the logarithmic ratio of fluorescence emission intensities at excitation wavelengths of 410 nm and 454 nm: log (*I*_410 nm_/ *I*_454 nm_). To construct the calibration curve, the fluorescence ratios were plotted against the pH of standard solutions (33 µM pyranine; 5 mM KH_2_PO_4_; 100 mM KCl) across a pH range of 5 to 7.5.

### Determination of membrane potential dynamics in *B. fragilis*

Wild-type and Δ*agcT* mutant *B. fragilis* cells were cultured under microaerobic conditions until reaching the logarithmic growth phase. The cultures were then diluted to an OD_600_ of 0.1 in growth medium supplemented with 0.5 mg mL^-1^ fatty acid-free bovine serum albumin (BSA). Cells (200 μl) were transferred to black polystyrene 96-well plates, and their autofluorescence was recorded for up to 5 minutes. DiSC_3_(5), dissolved in DMSO, was subsequently added to a final concentration of 2 μM (1% DMSO), and fluorescence quenching was monitored until a stable baseline was established. Fluorescence intensity was measured every minute, with vigorous shaking between readouts, using an excitation wavelength of 651 nm and an emission wavelength of 675 nm. All media, plates, and instruments were pre-warmed to 37°C before use.

### Determination of the activity of electron transport chain complexes

To assess the activity of respiratory complexes, inside-out membrane vesicles were prepared from *Bacteroides fragilis* strains, including wildtype, Δ*agcT*, and the indicated mutants. Briefly, bacterial cells were converted into spheroplasts by incubation in 200 mM Tris-HCl (pH 8.0), 2 mM EDTA, and 30% sucrose containing 0.1 mg mL^-1^ lysozyme. Following centrifugation at 5,000 × *g* for 10 minutes, the spheroplasts were resuspended in 100 mM HEPES-KOH (pH 7.5), 50 mM KCl, 10 mM MgSO_4_, 2 mM dithiothreitol (DTT), and 0.5 mM phenylmethanesulfonyl fluoride. The suspension was sonicated, and cell debris was removed by centrifugation at 5,000 × *g* for 10 minutes. Membrane vesicles were collected by ultracentrifugation at 120,000 × *g* for 30 minutes and resuspended in 25 mM HEPES-KOH buffer (pH 7.5).

Cytochrome oxidase activity was quantified by monitoring the oxidation of N,N,N′,N′-tetramethyl-p-phenylenediamine (TMPD) in the presence of 0.2 mM ascorbate. Absorbance at 611 nm was measured every 10 seconds over 10 minutes using 5 μg of membrane protein in a reaction containing 5 mM TMPD. NADH dehydrogenase activity was assessed via the reduction of 2,6-dichlorophenolindophenol (DCIP) at 600 nm, with absorbance measured over a 30-minute interval. Reactions contained 50 μM DCIP, 125 μM NADH, and 1 μg of membrane vesicles. Enzyme activity was expressed as absorbance change per minute per microgram of membrane protein.

### Statistical Analysis

Statistical analyses were performed using GraphPad Prism 10 and R v4.4.1 unless otherwise noted. For comparisons between two groups, two-tailed unpaired Student’s t-tests were used. For multiple group comparisons, one-way or two-way analysis of variance (ANOVA) was applied, followed by Tukey’s post hoc test where appropriate. For transposon sequencing (Tn-seq) data, gene fitness in the adult gut was assessed using the Wald test within the DESeq2 framework, with multiple testing correction using the Benjamini–Hochberg procedure. In the neonatal gut, due to sparse counts, fitness estimation was performed using a zero-inflated negative binomial (ZINB) model, and significance was determined by Wald z-test. All statistical tests were two-tailed unless specified, and significance thresholds were defined as follows: ^*^*p*-value < 0.05, ^**^*p*-value < 0.01, ^***^*p*-value < 0.001. P-values greater than 0.05 were annotated.

## Acknowledgments

We acknowledge the Gnotobiotic Core Facility at Harvard Medical School and staff for gnotobiotic animal resources and support. This work was supported in part by the National Institutes of Health (R01-AT010268 and R01-AI165987: S.F.O.) and National Research Foundation of Korea (Sejong Science Fellowship Grants RS-2024-00348702: K.H. and Nurturing Next-generation Researchers Fellowship RS-2024-00411992, B.G.). CD1d tetramers were provided by the NIH Tetramer Core Facility (contract number 75N93020D00005). We thank Dr. Meng Wu (Washington University in St. Louis) for helpful discussion.

## Author contributions

S.F.O and K.H. conceived the study, and D.L.K. contributed to study discussions. S.F.O. supervised the overall project and K.H. performed all experiments and analyzed the data. D.-J.J. performed and supported animal experiments and flow cytometry analysis. J.-S.Y. assisted with *in vitro* bacterial experiments. B.G. supported LC-MS analysis. K.H. and S.F.O. wrote the manuscript, and all authors contributed to the relevant discussion.

## Competing interests

S.F.O. and D.L.K. hold intellectual properties (patent and patent application) on BfaGCs.

## Notes

### Competing Interest Statement

S.F.O. and D.L.K. hold intellectual properties (patent and patent application) on BfaGCs (US patent 10,329,315).

