## Supplementary figures 1-10 for "Bacterial glycosphingolipids orchestrate colonization and immune modulation in neonatal host"

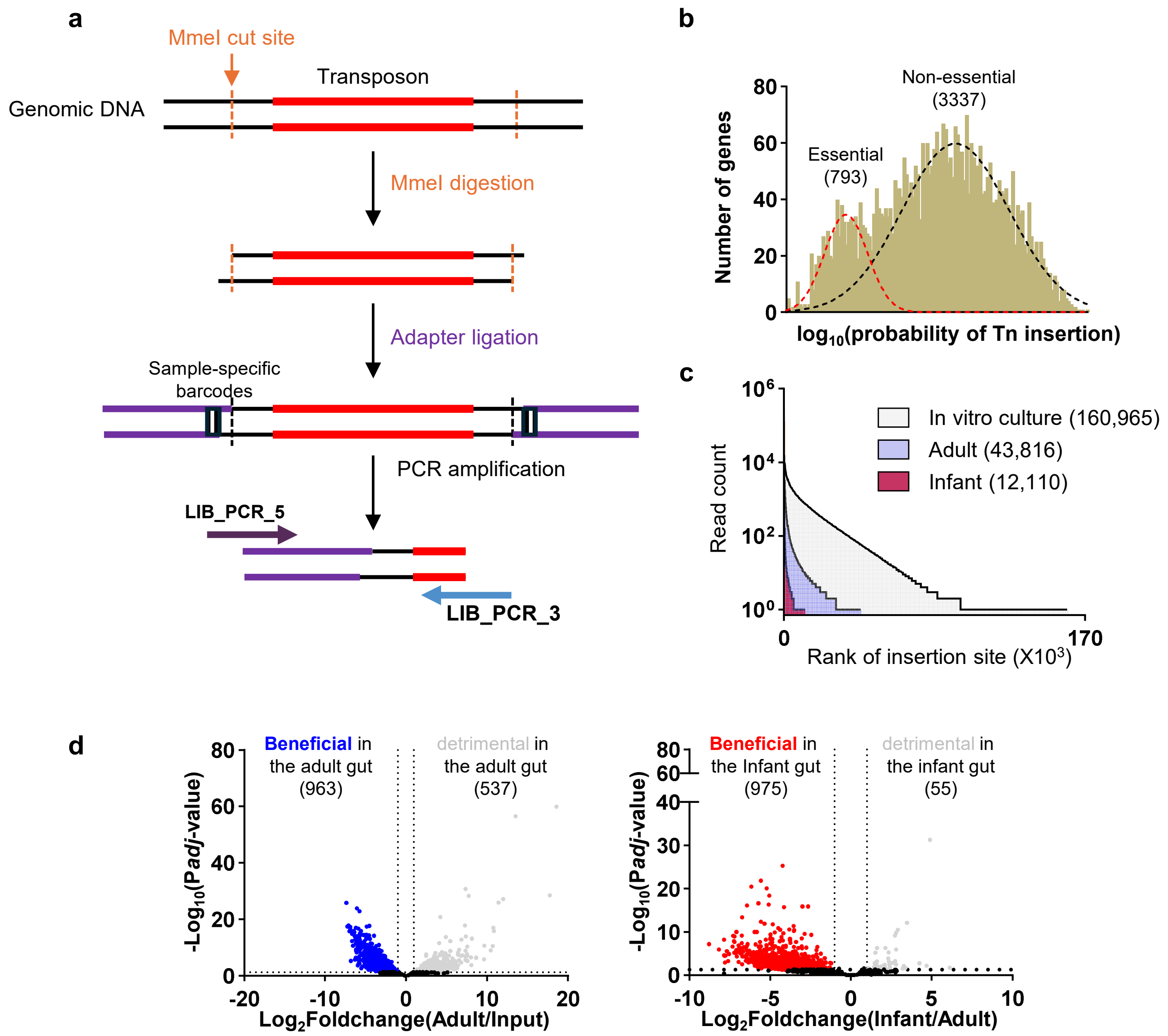


**Supplementary figure 1. Transposon sequencing library preparation and genome-wide fitness analysis**

**a,** Workflow for INSeq library preparation. Genomic DNA extracted from fecal samples was digested with MmeI, ligated to adapters containing sample-specific barcodes, and amplified by PCR to generate the sequencing library. **b,** Probability distribution of transposon (Tn) insertion across individual genes in the in vitro culture library pool. A two-component Gaussian mixture model was fitted to the bimodal distribution to distinguish essential (red) and non-essential (black) genes based on Tn insertion probabilities. **c,** Rank-ordered plot comparing Tn insertion read distributions in the in vitro pool (gray), adult (blue), and infant (red) samples. The number of unique insertion sites per groups is shown. Total read counts are as follows: In vitro culture, 40,656,127; Adult, 9,953,646; Infant, 1,615,306. **d,** Volcano plot illustrating genome-wide differences in bacterial gene fitness between in vitro culture and the adult gut (left) and between the adult gut and infant gut (right).

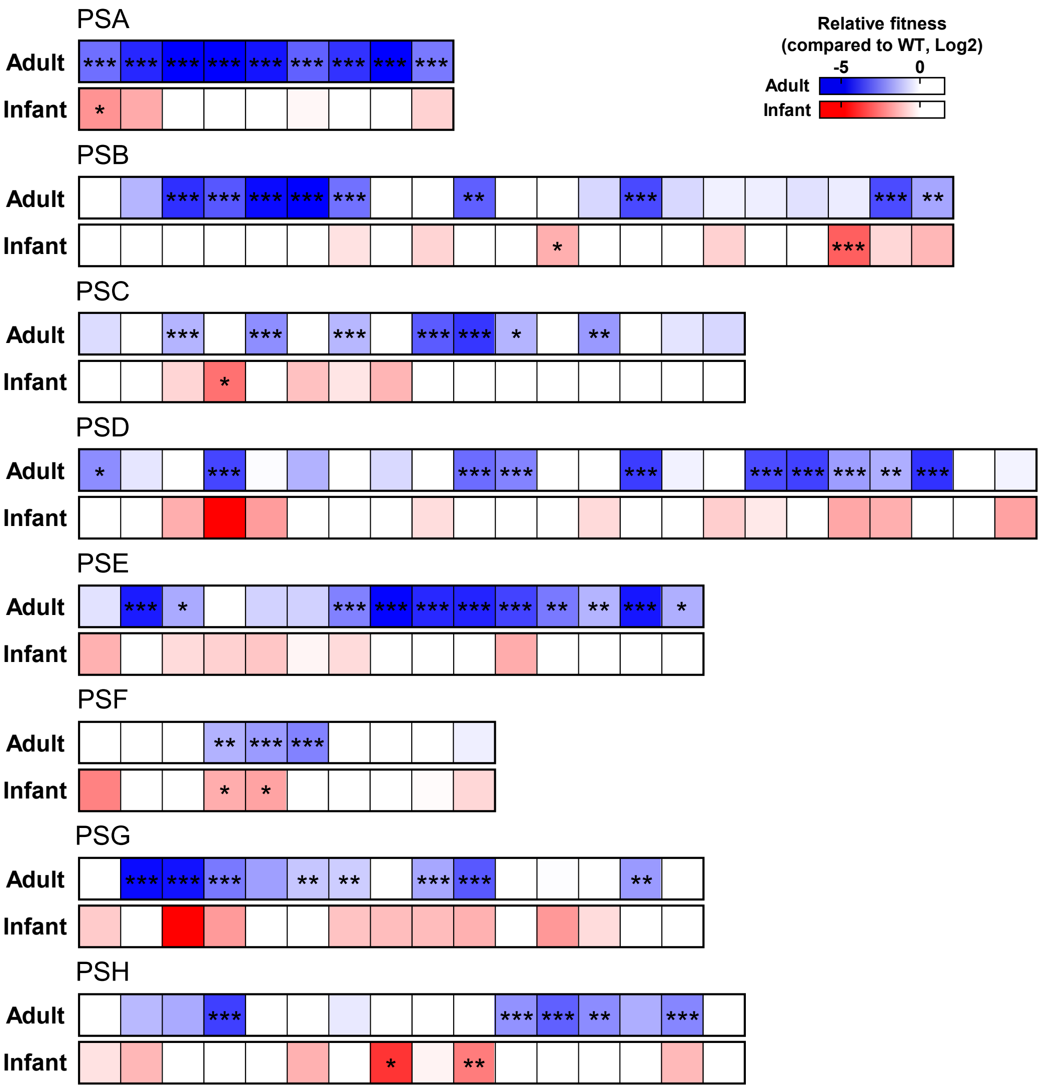


**Supplementary figure 2. Relative fitness of polysaccharide synthesis pathway mutants in the adult and infant gut**

The relative fitness of mutants for all genes involved in eight polysaccharide synthesis pathways (Polysaccharide A–H) was assessed in the adult and infant gut compared to the wild-type (WT) using Tn-seq analysis.

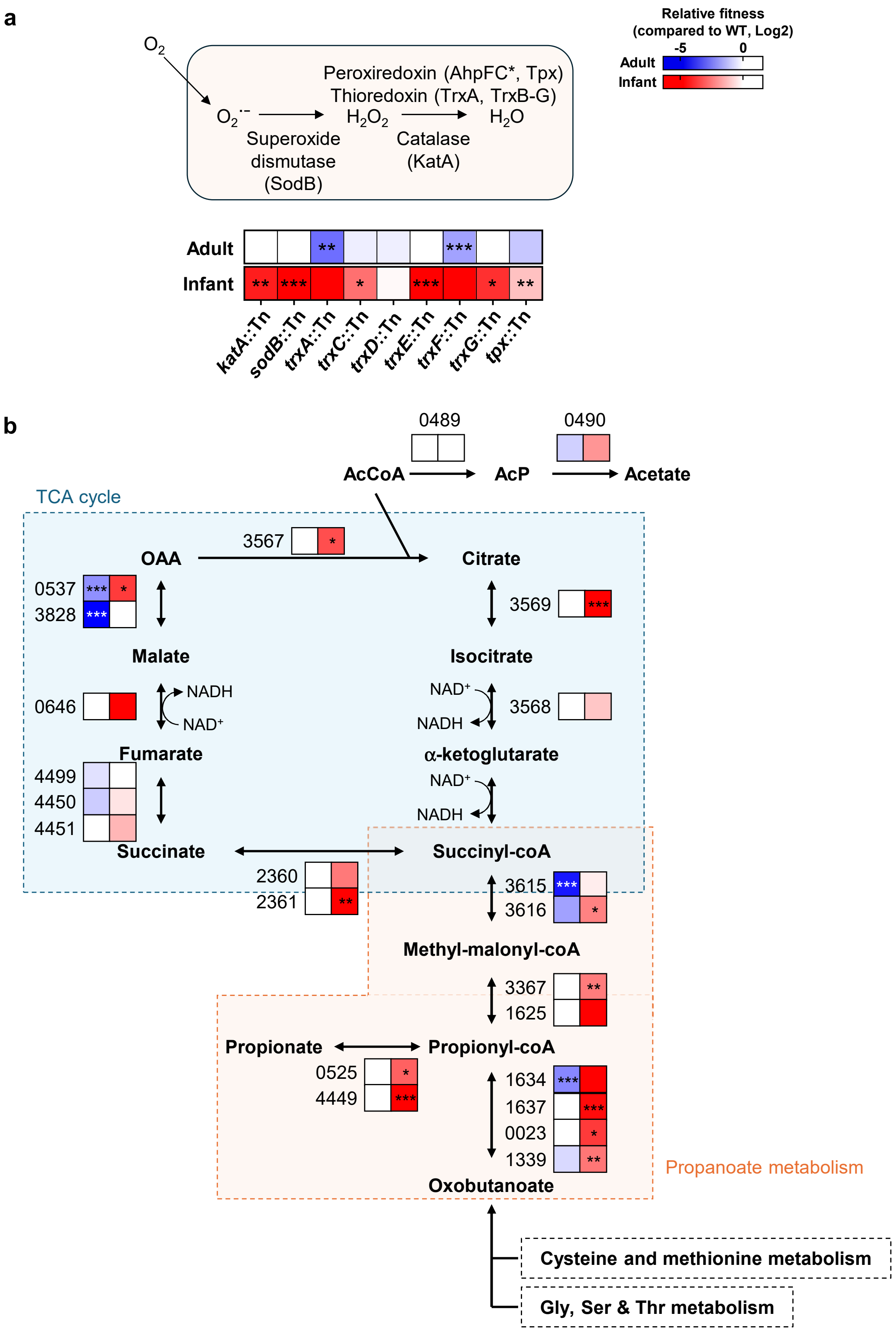


**Supplementary figure 3. Relative fitness of metabolic pathway mutants in the adult and infant gut**

The relative fitness of mutants for genes involved in a) reactive oxygen species (ROS)-scavenging and b) the TCA cycle, propanoate metabolism, and acetate synthesis was assessed in the adult and infant gut compared to the wild-type (WT) using Tn-seq analysis.

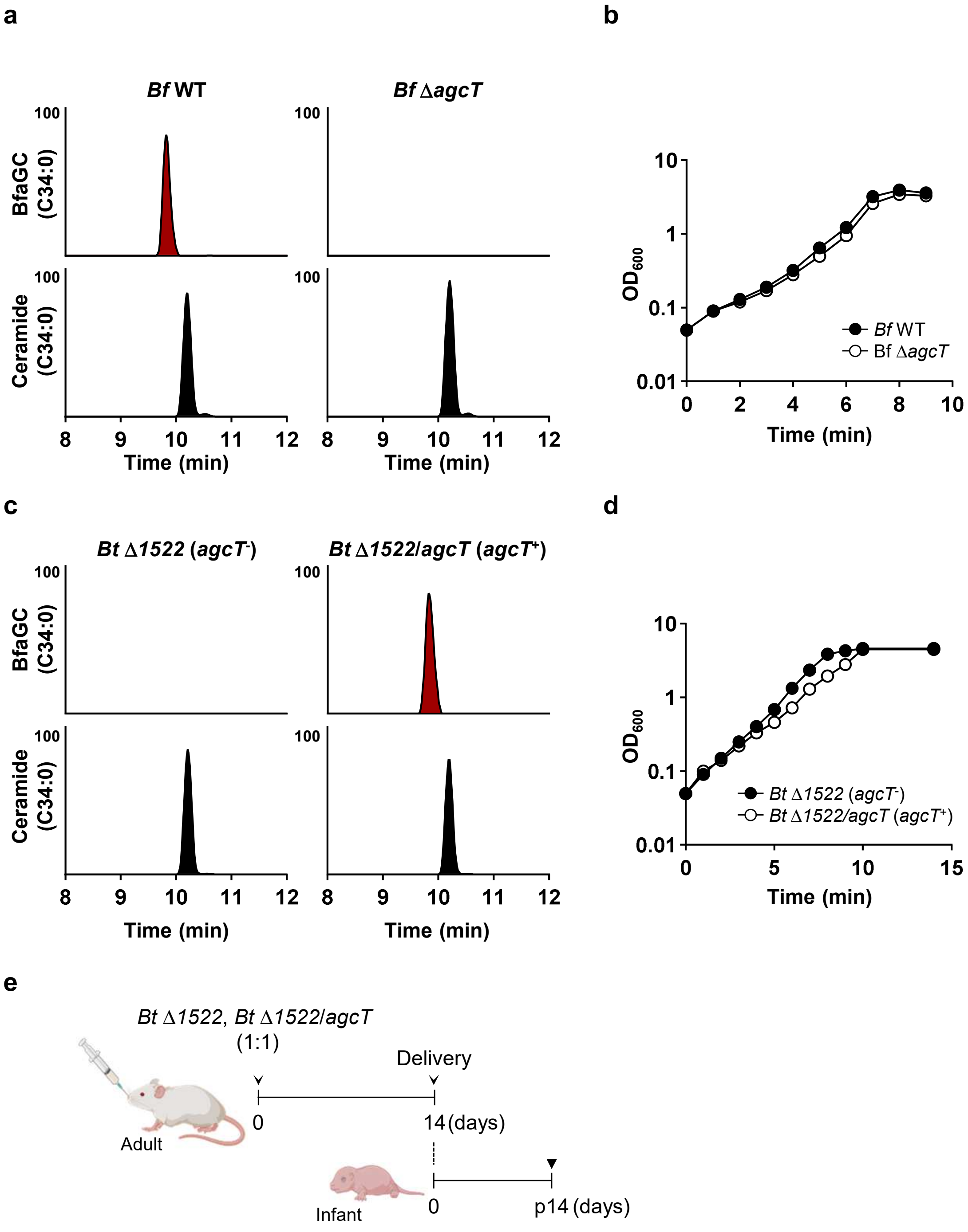


**Supplementary figure 4. Lipidomic profiling and growth analysis of aGC-producing Bacteroides species**

**a, c,** Relative abundance of *B. fragilis*-derived α-galactosylceramide (BfaGC, C34) and its precursor ceramide (C34) in **(a)** wild-type *B. fragilis* and its isogenic Δ*agcT* mutant, and **(c)** phosphoinositol dihydroxylceramide (PI-cer)-deficient *B. thetaiotaomicron* (Δ*1522*) strains either lacking (*agcT*$^{-}$) or expressing (*agcT*$^{+}$) the *agcT* gene. **b, d,** Growth curves of **(b)** wild-type and Δ*agcT B. fragilis*, and (d) PI-cer-deficient *B. thetaiotaomicron* (Δ*1522*) strains either lacking or expressing *agcT*, grown in PYG medium. **e,** Experimental design of co-colonization assays comparing the fitness of *B. thetaiotaomicron* strains with or without *agcT* expression.

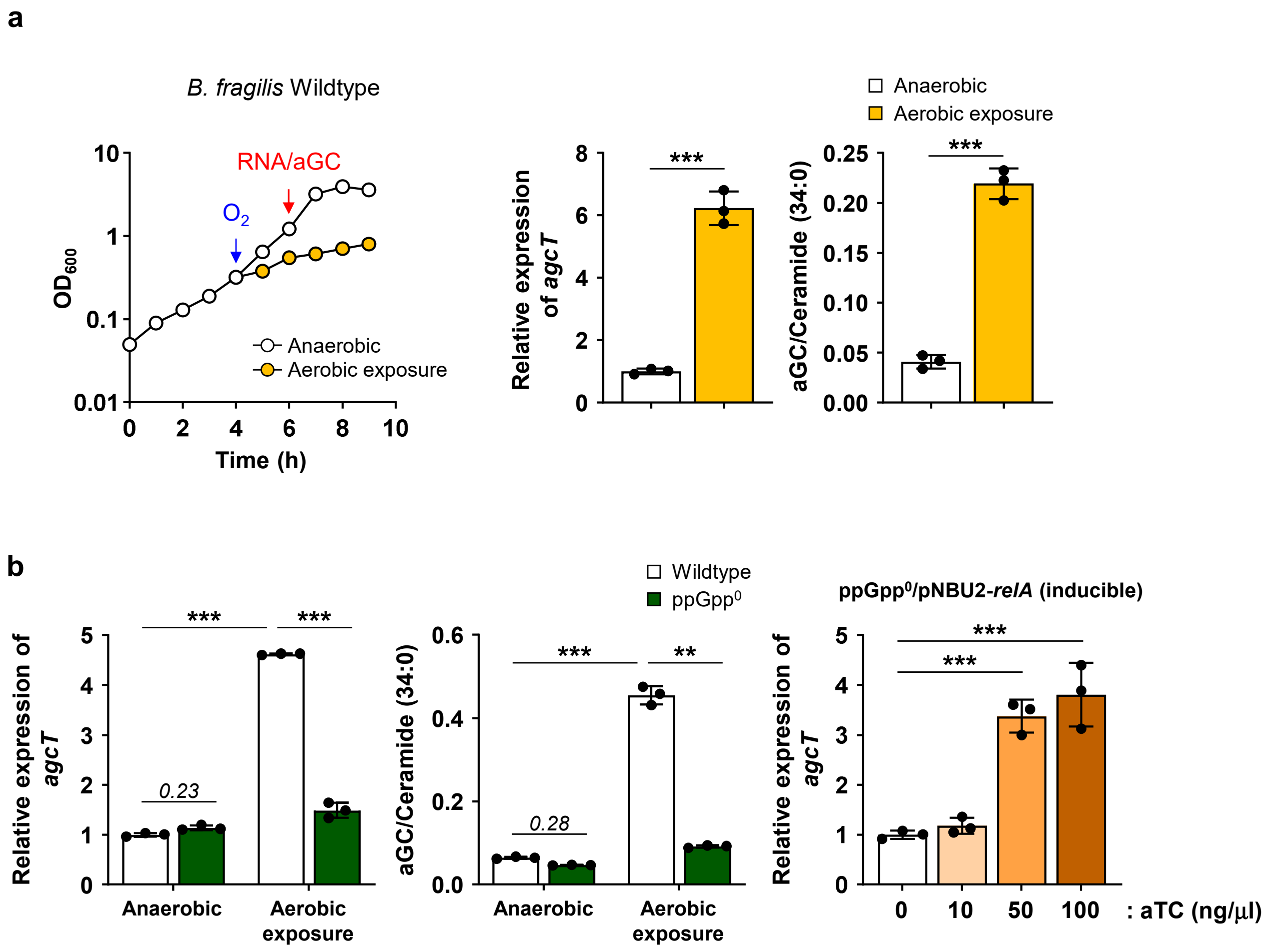


**Supplementary figure 5. Oxygen-dependent regulation of *agcT* expression and BfaGC synthesis in *B. fragilis***

**a,** Growth of wild-type *B. fragilis* in PYG medium under anaerobic and microaerobic conditions. During the exponential phase (OD ~0.3), half of the culture was exposed to microaerobic conditions, and growth was monitored. Two hours after oxygen exposure, cells were harvested for lipidomic and transcriptomic analysis. **b,** The expression of *agcT* and synthesis of α-galactosylceramide (BfaGC) in wild-type *B. fragilis* cultured under anaerobic and microaerobic conditions, measured by qRT-PCR and LC-MS, respectively. **c,** The expression of *agcT* and synthesis of BfaGC in wild-type *B. fragilis* and its isogenic ppGpp$^0$ mutant (Δ*0658*, *2136*) under anaerobic and microaerobic conditions, measured by qRT-PCR and LC-MS. **d,** The *agcT* expression in the ppGpp$^0$ mutant carrying the pNBU2-relA vector, cultured in the presence of the indicated concentrations of the inducer anhydrotetracycline.

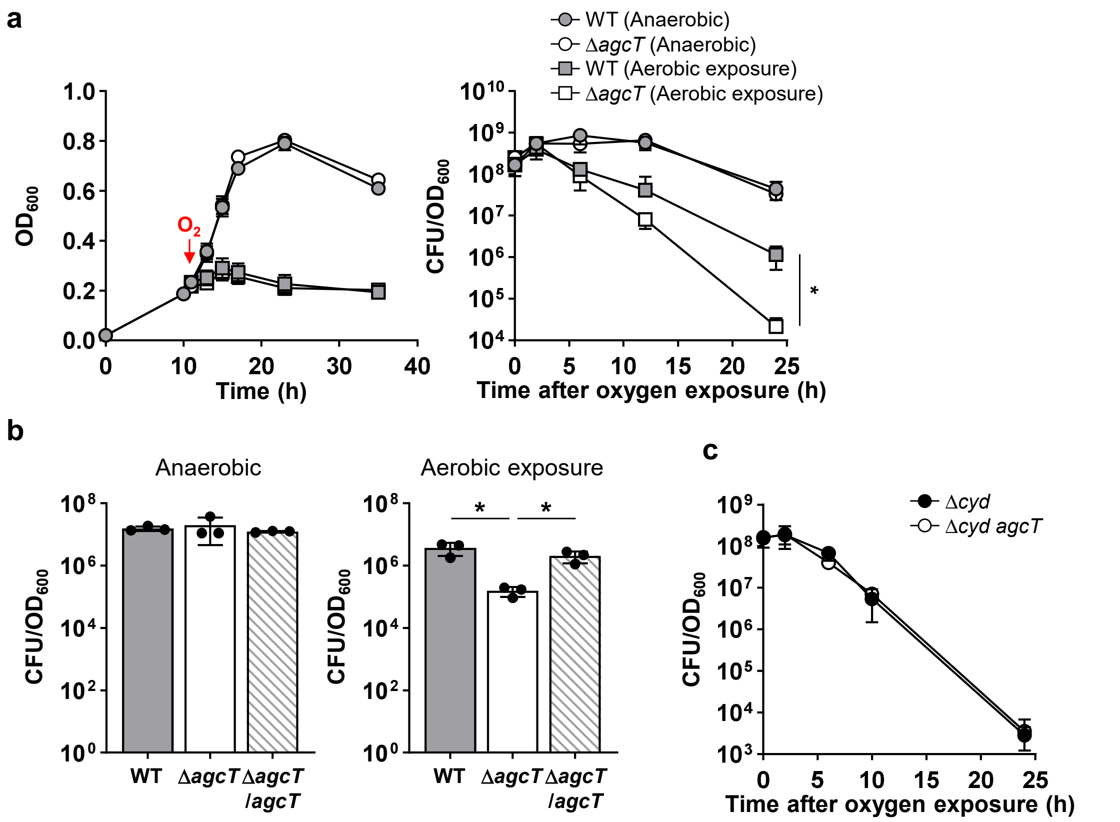


**Supplementary figure 6. BfaGC enhances aerotolerance under microaerobic condition**

**a,** Viability assay showing that the Δ*agcT* mutant exhibits reduced survival following aerobic exposure compared to WT, despite similar growth rates under both conditions. Viability was calculated by normalizing colony-forming units (CFUs) to OD600. **b,** Complementary expression of *agcT* restores the aerobic survival defect of the Δ*agcT* mutant. **c,** Viability of Δ*cyd* and Δ*cyd agcT* mutants after oxygen exposure.

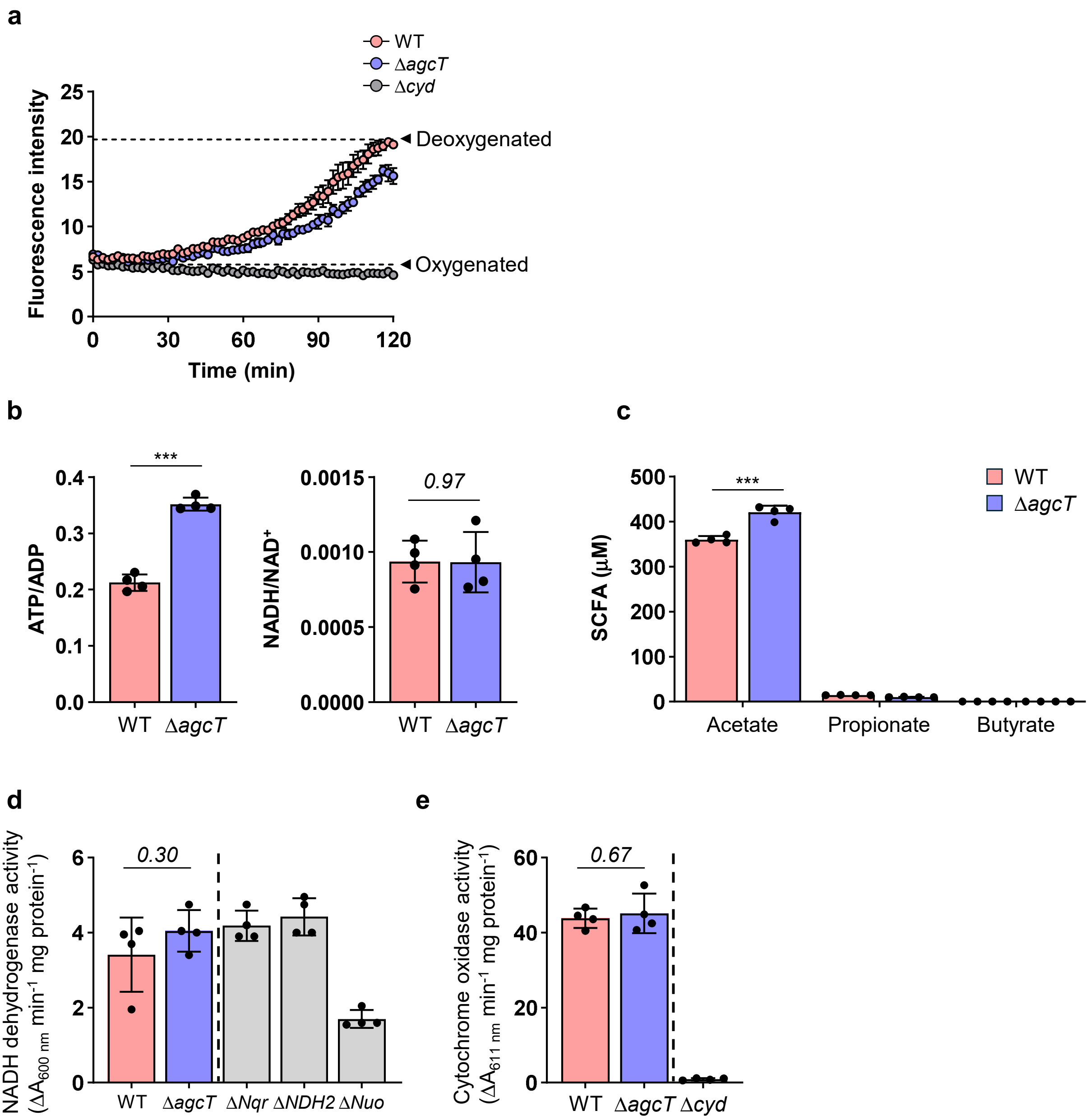


**Supplementary figure 7. BfaGC supports aerobic respiration under oxygen exposure**

**a,** Oxygen consumption rate of WT, Δ*agcT*, and Δ*cyd B. fragilis* strains measured under microaerobic conditions at 37°C. Oxygen depletion was inferred from increased fluorescence over a 2-hour period. values were normalized to dye-free controls. Cell-free oxygenated and deoxygenated media were included as baselines. **b,** Relative ATP/ADP and $NADH/NAD^+$ ratios, calculated from LC-MS signal intensities in WT and Δ*agcT* strains cultured under microaerobic conditions. **c,** Concentration of short-chain fatty acids (SCFAs) in culture supernatants collected 2 hours post-incubation under microaerobic conditions, measured by LC-MS following derivatization with 3-nitrophenylhydrazine. **d-e,** Activity of electron transport chain (ETC) complexes in inside-out membrane vesicles of *B. fragilis*. **d,** NADH dehydrogenase activity assessed via DCIP reduction (absorbance at 600 nm; 1 μg protein). **e,** Cytochrome oxidase activity measured by TMPD oxidation (absorbance at 610 nm; 5 μg protein).

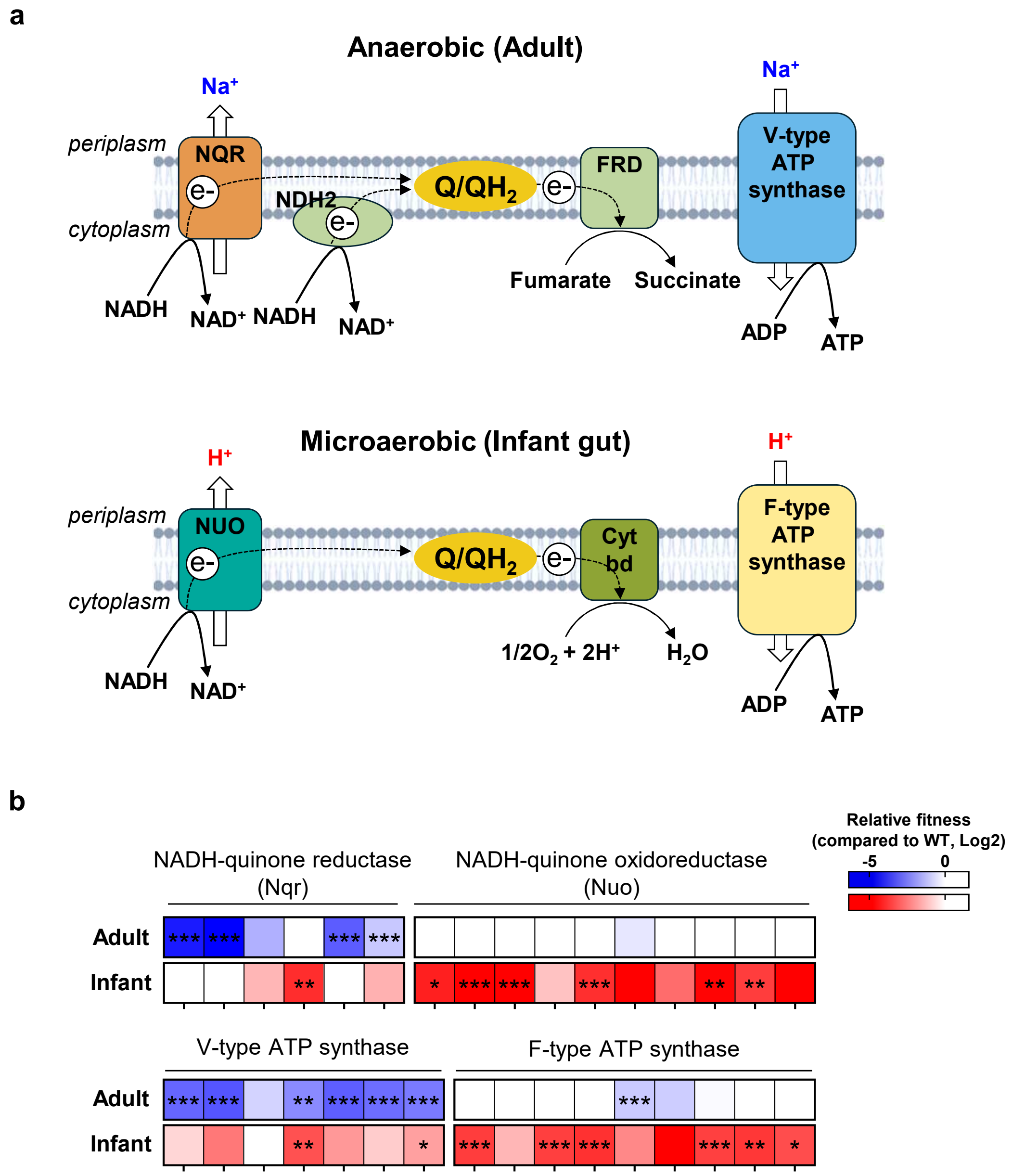


**Supplementary figure 8. Distinct electron transport chain configurations of *B. fragilis* in the adult and infant gut**

**a,** Graphical model of the distinct electron transport chain (ETC) configurations of *B. fragilis* in the adult and infant gut, established based on Tn-seq results. In the anaerobic adult gut, Nqr and NDH2 function as NADH dehydrogenases, transferring electrons to the quinone pool, which subsequently donates electrons to fumarate via fumarate reductase (FRD) to generate succinate. During this process, Nqr pumps sodium ions into the periplasmic space, generating a sodium motive force, which is utilized by V-type ATPase for ATP synthesis. In the microaerobic infant gut, Nuo functions as the NADH dehydrogenase, transferring electrons to the quinone pool and ultimately to oxygen via cytochrome *bd*. This process generates a proton motive force across the inner membrane, which is used by F-type ATP synthase to produce ATP. **b,** The relative fitness of mutants for all component genes of two NADH dehydrogenases (Nqr and Nuo) and two ATP synthases (V-type and F-type) was assessed in the adult and infant gut compared to the wild-type (WT) using Tn-seq analysis.

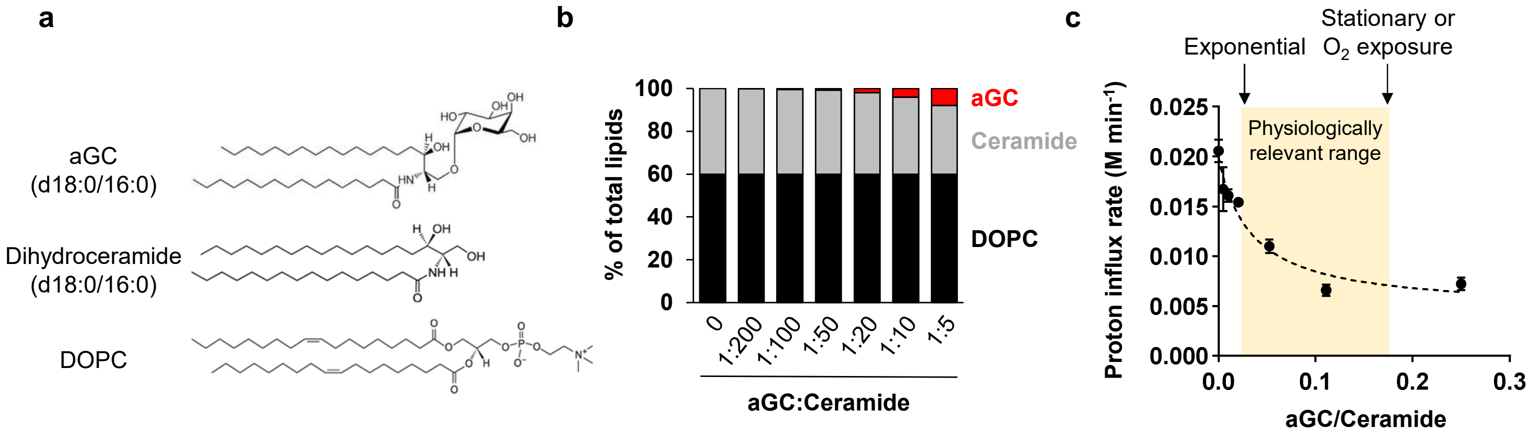


**Supplementary figure 9. BfaGC reduces proton permeability in synthetic lipid vesicles.** **a,** Chemical structures of synthetic lipids used for liposome generation. aGC, alpha-galactosylceramide; DOPC, 1,2-dioleoyl-sn-glycero-3-phosphatidylcholine. **b,** Liposome composition: 60% DOPC and 40% mixture of aGC and dihydroceramide at the indicated ratios. **c,** Proton permeability was assessed using small unilamellar vesicles (SUVs) composed of these synthetic lipids. SUVs were exposed to an external pH gradient (neutral to acidic), and intravesicular pH changes were monitored using the ratiometric fluorescent dye pyranine. Proton influx was quantified based on pH decay kinetics. The tested aGC/ceramide ratios reflect physiological levels observed in *B. fragilis* across growth phases (0.02 in exponential phase and up to 0.18 in stationary phase or under oxygen-exposed condition).

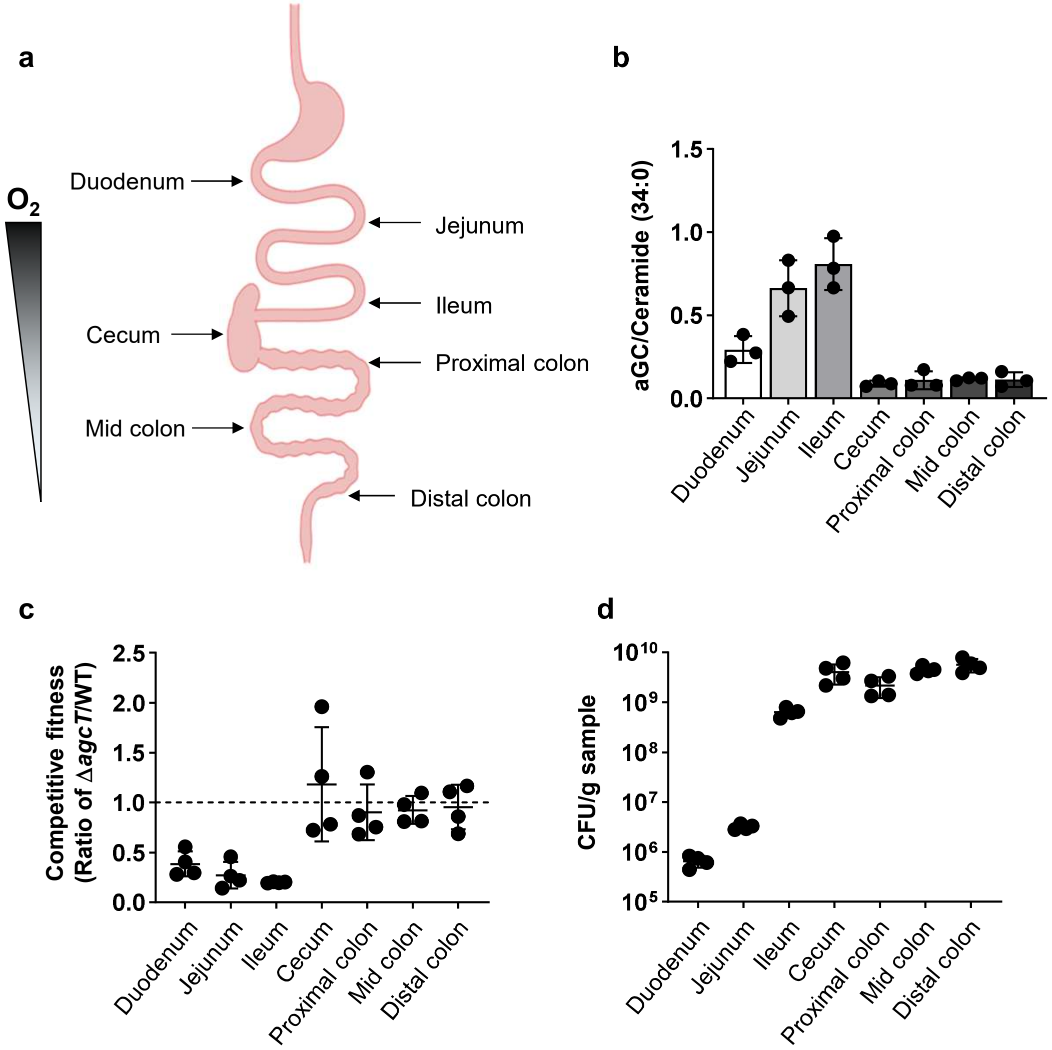


**Supplementary figure 10. Regional lipid composition and *B. fragilis* colonization along the gastrointestinal tract**

**a,** Graphical representation of the murine gastrointestinal (GI) tract, where oxygen levels decrease from the duodenum to the distal colon. Lipidomic profiling and colonization efficiency were assessed at the indicated regions (duodenum, jejunum, ileum, cecum, proximal colon, mid-colon, and distal colon). **b,** The proportion of α-galactosylceramide (aGC) to ceramide (C34) was measured by LC-MS analysis of samples collected from seven regions of the GI tract in *B. fragilis*-monocolonized mice. **c,** The competitive fitness of the *agcT* gene across the seven regions was determined through co-colonization experiments with wild-type *B. fragilis* and its isogenic Δ*agcT* mutant. **d,** Total *B. fragilis* colony-forming units (CFUs) were quantified in each of the seven regions in co-colonized mice.
